# High-density microbioreactor process designed for automated point-of-care manufacturing of CAR T cells

**DOI:** 10.1101/2023.04.07.535939

**Authors:** Wei-Xiang Sin, Narendra Suhas Jagannathan, Denise Bei Lin Teo, Faris Kairi, Dedy Sandikin, Ka-Wai Cheung, Yie Hou Lee, Rajeev J. Ram, Lisa Tucker-Kellogg, Michael E. Birnbaum

## Abstract

While adoptive cell therapies have revolutionized cancer immunotherapy, current autologous chimeric antigen receptor (CAR) T cell manufacturing face challenges in scaling to meet patient demands. CAR T cell production still largely rely on fed-batch, manual, open processes that lack environmental monitoring and control, whereas most perfusion-based, automated, closed-system bioreactors currently suffer from large footprints and working volumes, thus hindering process development and scaling-out. Here, we present a means of conducting anti-CD19 CAR T cell culture-on-a-chip. We show that T cells can be activated, transduced, and expanded to densities exceeding 150 million cells/mL in a two-milliliter perfusion-capable microfluidic bioreactor, thus enabling the production of CAR T cells at clinical dose levels in a small footprint. Key functional attributes such as exhaustion phenotype and cytolytic function were comparable to T cells generated in a gas-permeable well. The process intensification and online analytics offered by the microbioreactor could facilitate high-throughput process optimization studies, as well as enable efficient scale-out of cell therapy manufacturing, while providing insights into the growth and metabolic state of the CAR T cells during *ex vivo* culture.

## Introduction

Adoptive cell therapies have revolutionized the landscape of cancer immunotherapy, especially in the treatment of hematological malignancies. Some of the first patients to receive autologous cell therapies have been in complete remission for more than ten years^1^. In this span of time, six cell therapy products have since been approved by the United States Food and Drug Administration for the third-line treatment of relapsed or refractory B cell lymphomas and leukemias, and of multiple myeloma^2^. Despite the clinical success of these immunotherapies, the manufacturing of autologous cell therapies remains a complex, multi-step process. Patient peripheral blood mononuclear cells (PBMCs) are collected by leukapheresis, with T cells then isolated, activated, and genetically modified to express a chimeric antigen receptor (CAR). The CAR T cells are then expanded to clinical dosages, which can take 7 to 14 days, before final formulation, cryopreservation and subsequent thawing for patient infusion^3, 4^. Current commercial CAR T cell therapies largely rely on a centralized manufacturing model, but low manufacturing throughputs and long turnaround times of 21 to 37 days have hindered patient accessibility and treatment efficacy^4–8^. There is thus a need to intensify the scaling-out of autologous CAR T cell production, in order to reduce the wait times and cost of goods of these living drugs^3, 9, 10^.

To address these manufacturing challenges, there are increasing efforts to better understand the bioprocess design space of T cell cultures according to Quality by Design (QbD) principles, and to move towards decentralized and automated point-of-care manufacturing of autologous cell therapies in closed systems^11–13^. Currently, research- and preclinical-scale production of CAR T cells are commonly performed in open systems, such as well plates, T flasks and gas-permeable culture vessels, which may be prone to microbial contamination^14, 15^. These are typically fed-batch cultures with many manual handling steps, which can lead to fluctuations in nutrient availability and waste elimination^14, 16^. Further, process and operator variabilities may confound process development and prove difficult to translate to larger culture systems^17, 18^. While the closed-system G-Rex culture vessels (Wilson Wolf) were developed to provide semi-automated liquid handling^19^, and the CliniMACS Prodigy (Miltenyi Biotec) was developed to automate many steps of the CAR T cell manufacturing workflow^20^, these devices are essentially still based on fed-batch cultures with periodic medium exchanges, and lack real-time sensing of process parameters such as dissolved O_2_ and pH levels. On the other hand, various bioreactors, which can provide online monitoring and control of temperature, CO_2_, dissolved O_2_, and pH levels, have been explored for T cell culture and expansion^16, 21, 22^, and a subset of these, such as rocking bag bioreactors^23^ and hollow-fiber bioreactors^24^, are capable of running in perfusion culture modes. However, these bioreactors typically have large footprints, operate at large working volumes (hundreds of milliliters to liter scale) and require high seeding cell numbers (tens to hundreds of millions of cells)^16, 21, 22^. These factors, coupled with limited or costly starting materials and Good Manufacturing Practice (GMP)-grade reagents, such as PBMCs, viral vectors, medium, serum, and cytokines, make these platforms not well suited for high-throughput process optimization and not amenable to efficient scale-out of autologous cell therapy production. There is thus a gap for cell therapy manufacturing and process development with scaled-down perfusion-capable microbioreactors^25^, which, due to their small working volumes and footprints, can potentially enable greater process intensification.

A two-milliliter perfusion-capable microfluidic bioreactor, recently commercialized as the Mobius Breez microbioreactor (Erbi Biosystems, MilliporeSigma), has been developed for cell culture-on-a-chip^26, 27^. The Breez incorporates integrated mixers, injectors, and sensors for real-time monitoring and closed-loop control of flow rate, optical density, temperature, CO_2_, dissolved O_2_, and pH levels. It has been used for both microbial and mammalian cell cultures, such as *E. coli* for plasmid DNA production^28^, *P. pastoris* for recombinant human growth hormone and interferon-alpha production^29, 30^, and Chinese Hamster Ovary (CHO) cells for monoclonal antibody production, for which viable cell densities were shown to be comparable to those achieved in a bench-scale stirred-tank bioreactor^31, 32^. Here, we established and assessed a CAR T cell production process on this microbioreactor, with T cell activation, lentiviral transduction, and CAR T cell expansion steps all happening within the microfluidic chip. Breez-based CAR T cell production achieved more than 200 million viable CAR+ T cells, which met the minimum cell dose of tisa-cel (Kymriah) and exceeded the maximum cell dose of axi-cel (Yescarta). Importantly, the anti-CD19 CAR T cells produced at high densities in the microbioreactor were highly functional as measured via cytokine secretion and cytotoxic activity when co-cultured with CD19+ tumor cells *in vitro*. Phenotypically, despite somewhat higher proportions of T cells in the microbioreactor expressing differentiation and senescence markers, exhaustion markers were comparable to T cells cultured in the gas-permeable well. Additionally, continuous sensing of the microbioreactor environment provided information complementary to the measurement of metabolite levels in spent media. From this data, computational modeling estimated donor-specific rates of expansion and metabolism, thus providing a non-destructive readout of cell health and state while ensuring sterility.

## Results

### Establishment of CAR T cell process on microbioreactor versus gas-permeable well plate

While the Breez has previously been demonstrated for the culture of mammalian cell lines^31, 32^, its use in any human primary cell culture, or in the production autologous cell therapy products, has not been previously reported. We thus set out to establish and benchmark CAR T cell production protocols suitable for use in a microbioreactor. The Breez microbioreactor is comprised of a base station controller and a CO_2_ controller supporting up to four “pods”, each of which can house a microfluidic chip linked to a bottle rack assembly, supplied as a sterile, single-use consumable (**Supplementary Figure 1A**). This modular design allows highly parallelized synchronous or asynchronous running of up to four separate cultures. The microfluidic chip or “cassette” houses the growth chamber with a 2 mL culture volume; integrated reservoirs, injectors, pumps, valves, and a cell-retention perfusion filter; as well as embedded optical density, dissolved O_2_, and pH sensors, which enable real-time monitoring and closed-loop control of these process parameters (**Figure 1A**). There is also an inoculation port for introducing cells or viral vectors into the growth chamber, as well as a sampling port for in-process sampling of cell suspension from the growth chamber (**Figure 1A**). The growth chamber within the plastic microfluidic chip is defined by a flexible, gas-permeable polydimethylsiloxane (PDMS) silicone membrane, the cyclical deflection of which enables fast, diffusion-based gas transfer and low-shear, bubble-free mixing of the growth chamber (**Figure 1B**). This cassette is connected to a bottle rack assembly, which can supply up to four fluid inputs, in addition to two fluid outputs – a cell-free perfusion output and a cell-containing waste output (**Supplementary Figure 1A**). In our experiments, three fluid inputs were used – culture medium for perfusion, water for evaporation compensation, and a basic carbonate/bicarbonate solution for pH adjustment. When using aseptic connections via tube welders and tube sealers, this closed-system microbioreactor can be wholly operated on the benchtop with reduced cleanroom requirements. For our proof-of-concept experiments, cell or viral vector inoculation and cell harvest were performed by connecting syringes to the inoculation port or the waste bottle, respectively, in a biosafety cabinet (BSC), with the rest of the process occurring outside of a BSC in an academic tissue culture room facility.

**Figure 1:**
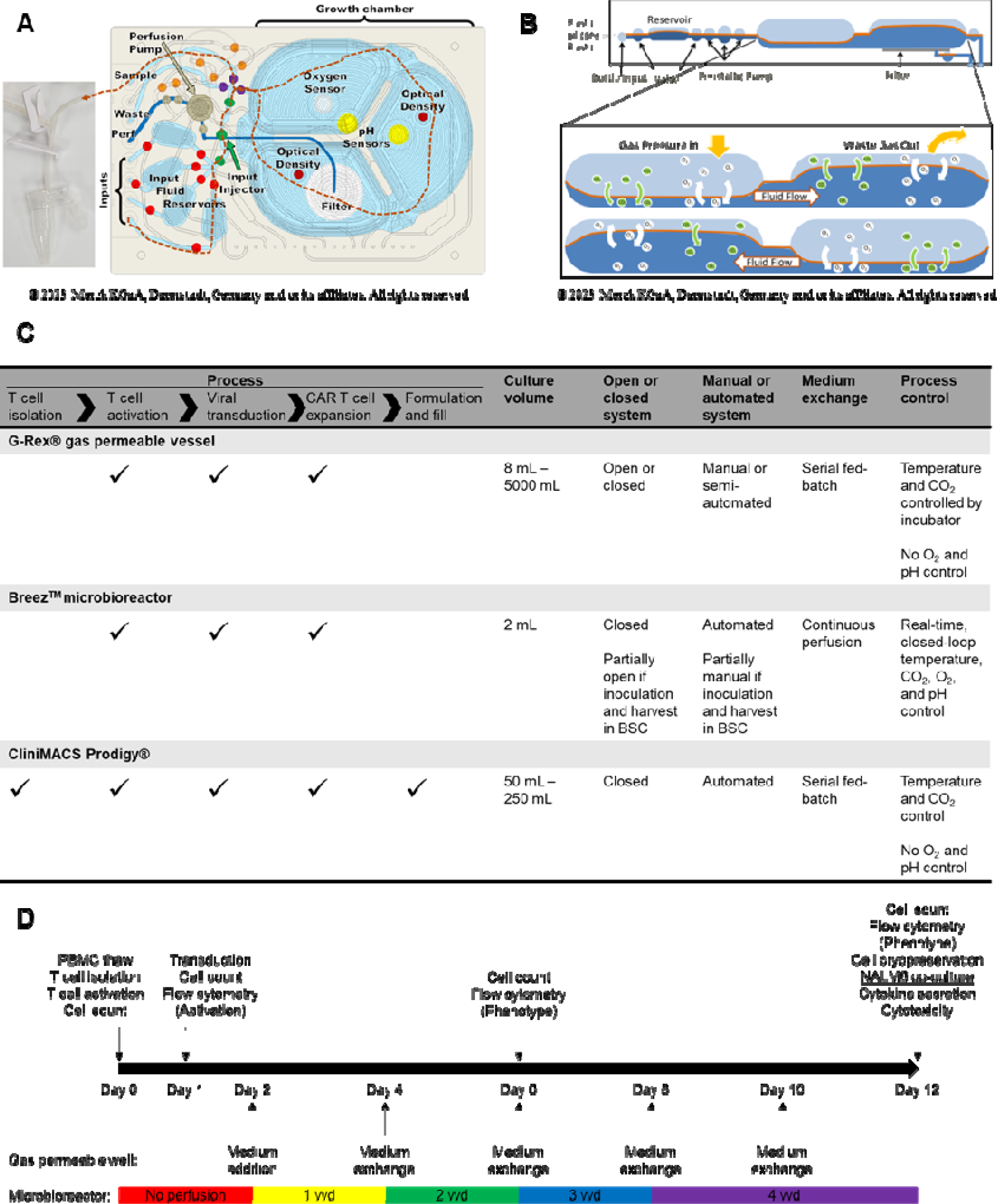
Schematics of CAR T cell production process on the microbioreactor and gas-permeable well plate. (A) Top view of the microbioreactor cassette, which contains the growth chamber with a 2 mL culture volume; integrated reservoirs, injectors, pumps, valves, and a cell-retention perfusion filter; as well as embedded optical density, dissolved O_2_, and pH sensors. (B) Side view of the microbioreactor cassette, illustrating a cross-section of the growth chamber defined by a flexible, gas-permeable polydimethylsiloxane (PDMS) silicone membrane, the cyclical deflection of which enables fast, diffusion-based gas transfer and low-shear, bubble-free mixing. (C) Comparison of T cell production process on the microbioreactor, gas-permeable wells, and CliniMACS Prodigy, and key features of the culture platforms. (D) Timeline of CAR T cell production process for the experiments described in this study. Medium exchange schedule on the gas-permeable wells, and perfusion flow rates for the microbioreactor are as indicated. At specific timepoints in the process, samples were taken for cell counts and flow cytometric analyses. Flow cytometry staining for activation markers was performed one day after activation, and that for surface phenotype markers was performed at the middle and end of the process. Fresh end-of-production anti-CD19 CAR T cells were co-cultured with CD19+ NALM6 cells for cytokine secretion assays. Cryopreserved end-of-production T cells were thawed and co-cultured with NALM6 cells for cytotoxicity assays. As biological replicates, PBMCs from three different healthy donors were used as starting materials, and CAR transduction was performed in technical replicates (at least three wells/cassettes per donor), with one non-transduced well/cassette as a control.

We compared the microbioreactor against a culture platform of the next smallest scale – a gas-permeable G-Rex 24-well plate with an 8 mL culture volume – via a 12-day CAR T cell production process, analogous to the CliniMACS Prodigy T Cell Transduction (TCT) Process, with T cell activation, lentiviral transduction, and CAR T cell expansion steps all happening within the respective culture vessels (**Figure 1C-D; Supplementary Figure 1B-C**). As biological replicates, PBMCs from three different healthy donors were used as starting materials, and CAR transduction was performed in technical replicates (at least three wells/cassettes per donor), with one non-transduced well/cassette as a control. At specific timepoints in the process, samples were taken for cell counts and flow cytometric analyses (**Figure 1D**). We ran the gas-permeable well plate culture as a perfusion-mimic by performing 6 mL medium exchanges every other day, whereas continuous perfusion mode was employed for the microbioreactor, starting at 1 vessel volume per day (vvd) approximately 24 hours after transduction, ramping up by 1 vvd every other day to reach a maximum flow rate of 4 vvd from Day 8 onwards. This feeding regime resulted in a total medium usage of 33 mL for each gas-permeable well, and 59 mL for each microbioreactor cassette (**Supplementary Figure 1D**). The gas-permeable well plate was placed in a standard 37°C, 5% CO_2_ incubator without any dissolved O_2_ and pH control, while the microbioreactor cultures were controlled at the following set-points: temperature of 37°C, minimum CO_2_ levels of 5%, minimum dissolved O_2_ levels of 80% air saturation, and a pH of 7.40 ± 0.05 (to reflect the physiological definition of acidosis and alkalosis) (**Supplementary Figure 1E**).

### T cell activation levels and transduction efficiencies were high and comparable between the microbioreactor and the gas-permeable well plate

In CAR T cell production, starting T cells must be activated to enable transduction by standard lentiviral vectors^33^. For activation, we mixed purified T cells with CD3/CD28 Dynabeads (Thermo Fisher Scientific) in a 25 mL tube, and then seeded the cell-bead mixture into four gas-permeable wells and at least four microbioreactor cassettes for each donor, at the same cell density, cell:bead ratio, and culture volume for both systems (**Supplementary Figure 1C, 2A**). To assess the level of T cell activation, we pooled cell samples taken from the replicate wells/cassettes (due to small starting cell numbers) on Day 1 before transduction, and performed flow cytometric analyses for the activation markers CD69 and CD25 (**Supplementary Figure 2B**). We observed that > 95% of the cells were either CD69+ or CD25+ (**Figure 2A**), indicating that the levels of T cell activation were similar between the two culture systems. Consistent with this, the expression of low-density lipoprotein receptor (LDL-R), which is the receptor for vesicular stomatitis virus G-protein (VSV-G) pseudotyped lentiviral vectors, and known to be upregulated upon T cell activation^33, 34^, was also comparable for T cells activated in the microbioreactor and the gas-permeable well plate (**Supplementary Figure 3A**).

**Figure 2:**
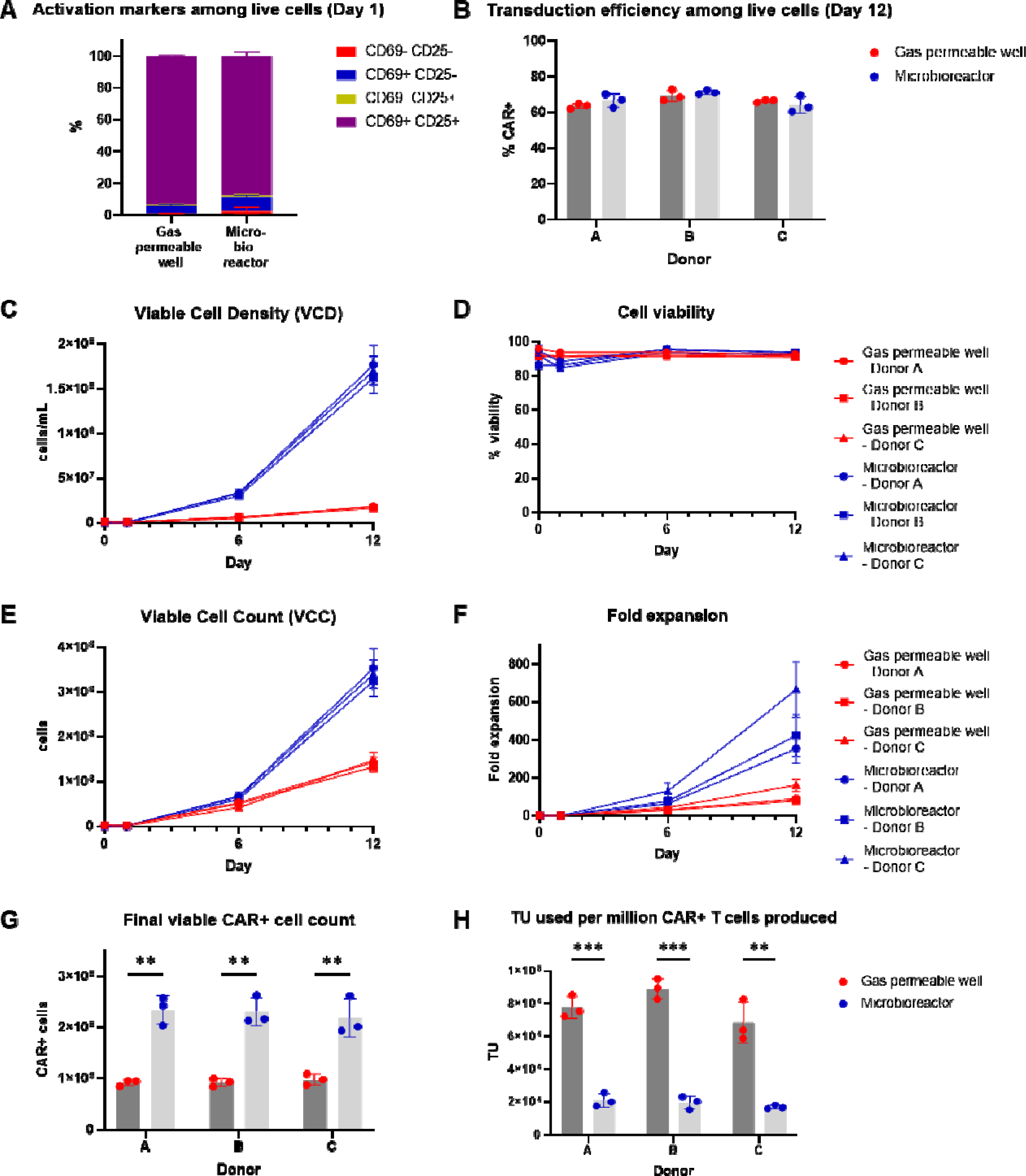
Comparable levels of activation and transduction, and significantly higher level of CAR T cell expansion on the microbioreactor compared with gas-permeable well plate. (A) Percentage of total live cells expressing the activation markers CD69 and/or CD25 on Day 1. Mean ± SD of n=3 donors. (B) Percentage of total live cells expressing CAR on Day 12. Mean ± SD of n=3 wells/cassettes. P > 0.05 by unpaired t test with Holm-Sidak’s multiple comparisons correction. (C) Viable cell density (VCD) in cells/mL over time, as determined by acridine orange-propidium iodide (AO-PI) staining. Mean ± SD of n=3 wells/cassettes. (D) Percentage cell viability as determined by AO-PI staining. Mean ± SD of n=3 wells/cassettes. (E) Viable cell count (VCC) in number of cells over time, as determined by AO-PI staining. Mean ± SD of n=3 wells/cassettes. (F) Fold expansion relative to VCC in cell sample taken after cell inoculation on Day 0, as determined by AO-PI staining. Mean ± SD of n=3 wells/cassettes. (G) Viable CAR+ cell count in number of CAR+ cells on Day 12, as determined by AO-PI staining. Mean ± SD of n=3 wells/cassettes. (H) Number of transduction units (TU) of lentiviral vector (LVV) used per million CAR+ cells produced. Mean ± SD of n=3 wells/cassettes. (G-H) ** P ≤ 0.01, *** P ≤ 0.001 by unpaired t test with Holm-Sidak’s multiple comparisons correction.

We next transduced cells with an anti-CD19 CAR-containing lentiviral vector (LVV) co-expressing GFP at multiplicity of infection (MOI) of 5. The total viable cell numbers were lower in the microbioreactor compared with the gas-permeable wells on Day 1 before transduction, hence lower amounts of LVV were required for transduction in the microbioreactor (**Supplementary Figure 3B**). For the gas-permeable wells, 1 mL of cell-free medium was replaced with 1 mL of fresh medium containing diluted LVV, and then left in static culture for 24 h before 6 mL of fresh medium was added to achieve the final culture volume of 8 mL (**Supplementary Figure 2A**). For the microbioreactor, 1 mL of cell-free medium was removed through the perfusion output, and then 1 mL of fresh medium containing diluted LVV was inoculated through the inoculation port. The growth chamber was continuously mixed for 2 h after the addition of LVV, and then subsequently reverted to intermittent mixing for the remainder of the process (**Supplementary Figure 2A**). We measured GFP expression as a proxy of CAR expression by flow cytometry on Days 6 and 12, and found that the CAR was stably expressed on T cells cultured on both culture systems (**Figure 2B; Supplementary Figure 3C-E**). The percentages of CAR+ T cells were largely comparable on total T cells between the microbioreactor and the gas-permeable well plate, ranging from 60% to 70% CAR+ (**Figure 2B; Supplementary Figure 3C-E**), indicating that both culture systems were equally capable of supporting high transduction efficiencies.

### Milliliter-scale perfusion-based microbioreactor enabled high-density production of CAR T cells at clinical dose levels

Having demonstrated equivalent levels of activation and transduction in the microbioreactor and the gas-permeable well plate, we next assessed the rates of CAR T cell expansion in the two culture systems. Compared with the gas-permeable wells, viable cell densities (VCD) in the microbioreactor were 5-fold higher on Day 6, and 10-fold higher on Day 12 (**Figure 2C, Supplementary Figure 4A**). Despite cell densities exceeding 150 million cells/mL in the microbioreactor, cell viabilities remained high and comparable to that in the gas-permeable wells (**Figure 2D, Supplementary Figure 4B**). Taking into account the 4-fold larger culture volume of the gas-permeable wells versus the microbioreactor, viable cell counts (VCC) in the microbioreactor were slightly higher on Day 6, and ∼ 2.5-fold higher on Day 12 (**Figure 2E, Supplementary Figure 4C**). These VCCs translate into a cumulative fold expansion exceeding 400-fold in the microbioreactor compared with less than 200-fold in the gas-permeable wells (**Figure 2F, Supplementary Figure 4D**), suggesting that improved medium exchange and/or gas transfer could possibly contribute to the better CAR T cell expansion in the microbioreactor. Notably, with a final VCC exceeding 300 million total T cells and 60-70% CAR+ cells, the final number of CAR+ viable T cells exceeded 200 million in the microbioreactor (**Figure 2G, Supplementary Figure 4E**), which met the minimum cell dose of tisa-cel (Kymriah) for pediatric/young adult B-cell acute lymphoblastic leukemia (ALL) and adult diffuse large B-cell lymphoma (DLBCL), and exceeded the maximum cell dose of axi-cel (Yescarta) for adult DLBCL^2, 35, 36^. Hence, a clinical dose of anti-CD19 CAR T cells can potentially be produced in a single microbioreactor cassette, thus substantially increasing the throughput and improving the capacity to scale-out autologous CAR T cell manufacturing for multiple patients in parallel.

Interestingly, the high CAR T cell numbers generated in the microbioreactor were achieved with significantly lower amounts of LVV used per million CAR+ T cells produced, compared with that used in the gas-permeable well plate (**Figure 2H**). This was due to lower amounts of LVV used at the start of production (**Supplementary Figure 3B**), and higher numbers of CAR+ T cells generated at the end of the process (**Figure 2G, Supplementary Figure 4E**). Finally, despite the perfusion-based microbioreactor requiring a higher total medium usage due to the continuous medium exchange compared with the serial fed-batch gas-permeable well plate (**Supplementary Figure 1D**), this was somewhat mitigated by the small-scale 2 mL culture volume, and compensated for by the significantly enhanced T cell expansion (**Figure 2F, Supplementary Figure 4D**), such that the medium efficiency of the perfusion-based microbioreactor process was comparable to or better than that of the perfusion-mimic gas-permeable well process (**Supplementary Figure 4F**).

### CAR T cells from both culture platforms were predominantly of Tn/Tscm and Tcm phenotype and of comparable exhaustion status

We next characterized the phenotype of the CAR T cells produced on the two culture systems (**Supplementary Figure 5**). High CD3+ purity was obtained after T cell isolation (**Supplementary Figure 6A**), and this was maintained through the end of the process for both culture systems (**Supplementary Figure 6B**). CD4+/CD8+ ratio decreased over time in culture, more significantly in the microbioreactor compared with the gas-permeable wells for Donors A and C, and regardless of culture system for Donor B (**Figure 3A**). Overall, the CD4+/CD8+ ratio was not drastically skewed in either culture system. We further analyzed the differentiation status of the CAR T cells produced on the two culture systems by flow cytometric staining of CCR7 and CD45RA. On Day 6, the percentage of cells in the naïve/stem cell memory (Tn/Tscm) (CCR7+ CD45RA+) compartment and the central memory (Tcm) (CCR7+ CD45RA–) compartment, subsets associated with longer *in vivo* persistence^37, 38^, was slightly lower in the microbioreactor compared with the gas-permeable wells (**Figure 3B**). While the percentage of Tn/Tscm and Tcm cells decreased over time in culture for both systems, the difference between the microbioreactor and the gas-permeable wells was also observed on Day 12, with the former having a slightly higher percentage of more differentiated effector memory (Tem) (CCR7– CD45RA–) and terminally differentiated effector (Temra) cells (CCR7– CD45RA+) (**Figure 3B**). The proportions of cells in the differentiation subsets were similar when gated on total T cells or on CAR+ T cells (**Supplementary Figure 6C-E**), indicating that CAR+ T cells were not phenotypically different from non-transduced T cells. In addition, the percentage of cells expressing CD127 (IL-7Rα), which has been reported to be upregulated on Tn and Tcm and downregulated on Tem^39^, were lower for cells expanded on the microbioreactor compared with the gas-permeable wells (**Supplementary Figure 6F**). Interestingly, we observed a donor-specific feature of Donor B, whose cells were substantially more differentiated at the end of the process compared with Donors A and C (**Figure 3B**), as well as another donor-specific feature of Donor C, whose cells had significantly higher expression of CD57, a marker typically associated with terminal effector or senescent T cells^40^, on both Day 6 and Day 12 in the microbioreactor compared with the gas-permeable wells (**Supplementary Figure 6G**). For Donors A and B, the percentage of CD57+ cells was higher in the microbioreactor on Day 12, but still below 5% of total T cells (**Supplementary Figure 6G**). Overall, differences in T cell differentiation status between the microbioreactor and the gas-permeable well plate were relatively subtle, and more than half of the end-of-production cells from both culture systems were of Tn/Tscm and Tcm phenotype, which have been recognized as a desirable attribute of CAR T cell therapies.

**Figure 3:**
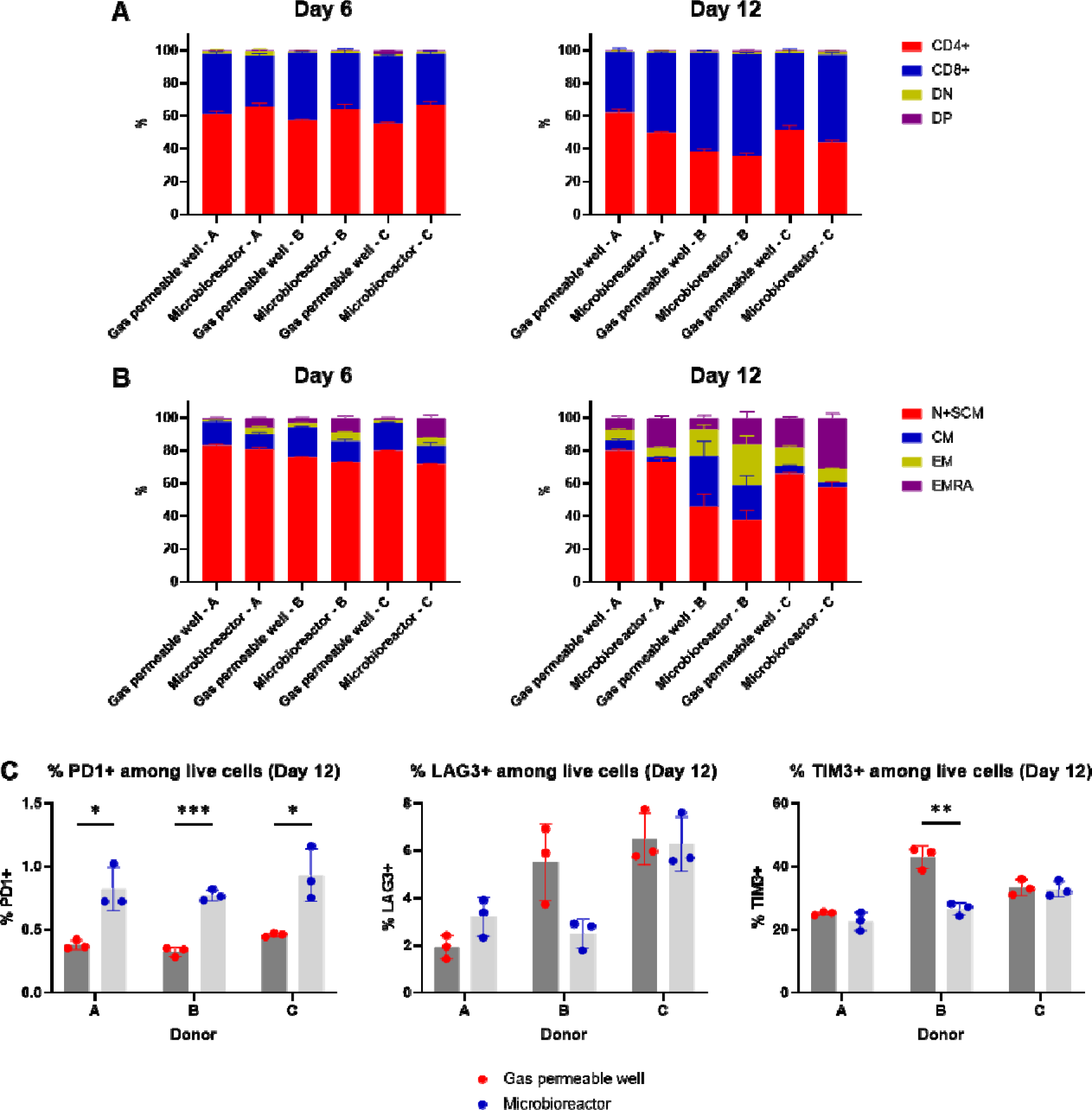
T cell differentiation and exhaustion phenotypes of CAR T cells from the microbioreactor and the gas-permeable well plate. (A) Percentage of total live cells that are CD4+, CD8+, CD4– CD8– (double negative, DN) and CD4+ CD8+ (double positive, DP) on Days 6 and 12. Mean ± SD of n=3 wells/cassettes. (B) Percentage of total live cells that are CCR7+ CD45RA+ (naïve and stem cell memory, Tn/Tscm), CCR7+ CD45RA– (central memory, Tcm), CCR7– CD45RA– (effector memory, Tem), and CCR7– CD45RA+ (terminally differentiated effector, Temra) on Days 6 and 12. Mean ± SD of n=3 wells/cassettes. (C) Percentage of total live cells expressing the exhaustion markers PD1, LAG3, and/or TIM3 on Day 12. Mean ± SD of n=3 wells/cassettes. * P ≤ 0.05, ** P ≤ 0.01, *** P ≤ 0.001 by unpaired t test with Holm-Sidak’s multiple comparisons correction.

T cell exhaustion has been reported to be detrimental to the effector function and persistence of CAR T cell therapies^41^. We looked at the T cell exhaustion markers PD-1, TIM-3, and LAG-3 on the CAR T cells generated from the two culture systems, and observed that the percentage of total T cells expressing PD-1 and LAG-3 was lower than 8%, with PD-1 being somewhat higher for cells expanded on the microbioreactor, and LAG-3 being comparable between the two culture systems (**Figure 3C**). The percentage of total T cells expressing TIM-3 was either comparable or lower for cells expanded on the microbioreactor compared with gas-permeable wells. Taken together, these data indicate that the T cell exhaustion levels were low and comparable between the microbioreactor and the gas-permeable well plate.

### CAR T cells generated on the microbioreactor were highly functional in cytokine secretion and cytolytic activity

Having looked at the phenotypic characteristics, we next evaluated the function of the CAR T cells that were produced at high densities on the microbioreactor. End-of-production anti-CD19 CAR T cells (and non-transduced T cells) were harvested from the microbioreactor and gas-permeable wells, and co-cultured with CD19+ NALM6 cells, following which cell culture supernatants were analyzed by Luminex multiplex assays. Out of 38 cytokines that were measurable in the NALM6 co-cultures (**Figure 4A**), 30 showed no significant differences, six were higher when the CAR T cells came from gas-permeable wells (G-CSF, GM-CSF, IL-3, PD-L1, TNF-L, and TNF-β) (**Supplementary Figure 7A, B, D-G**), and two were higher when the CAR T cells came from the microbioreactor (IL-5 and IL-13) (**Supplementary Figure 7I, J**). Additionally, while not statistically significant, gas-permeable well CAR T cells secreted higher levels of IFN-γ (**Supplementary Figure 7C**), whereas microbioreactor CAR T cells secreted higher levels of IL-4 (**Supplementary Figure 7H**). These data suggest that gas-permeable well CAR T cells secrete higher levels of pro-inflammatory and neurotoxicity-or cytokine release syndrome (CRS)-associated cytokines (GM-CSF, IFN-γ, and TNF-L) upon encountering tumor cells, while microbioreactor CAR T cells secreted higher levels of type 2 cytokines (IL-4, IL-5, and IL-13), which have been associated with long-term remission^42^.

**Figure 4:**
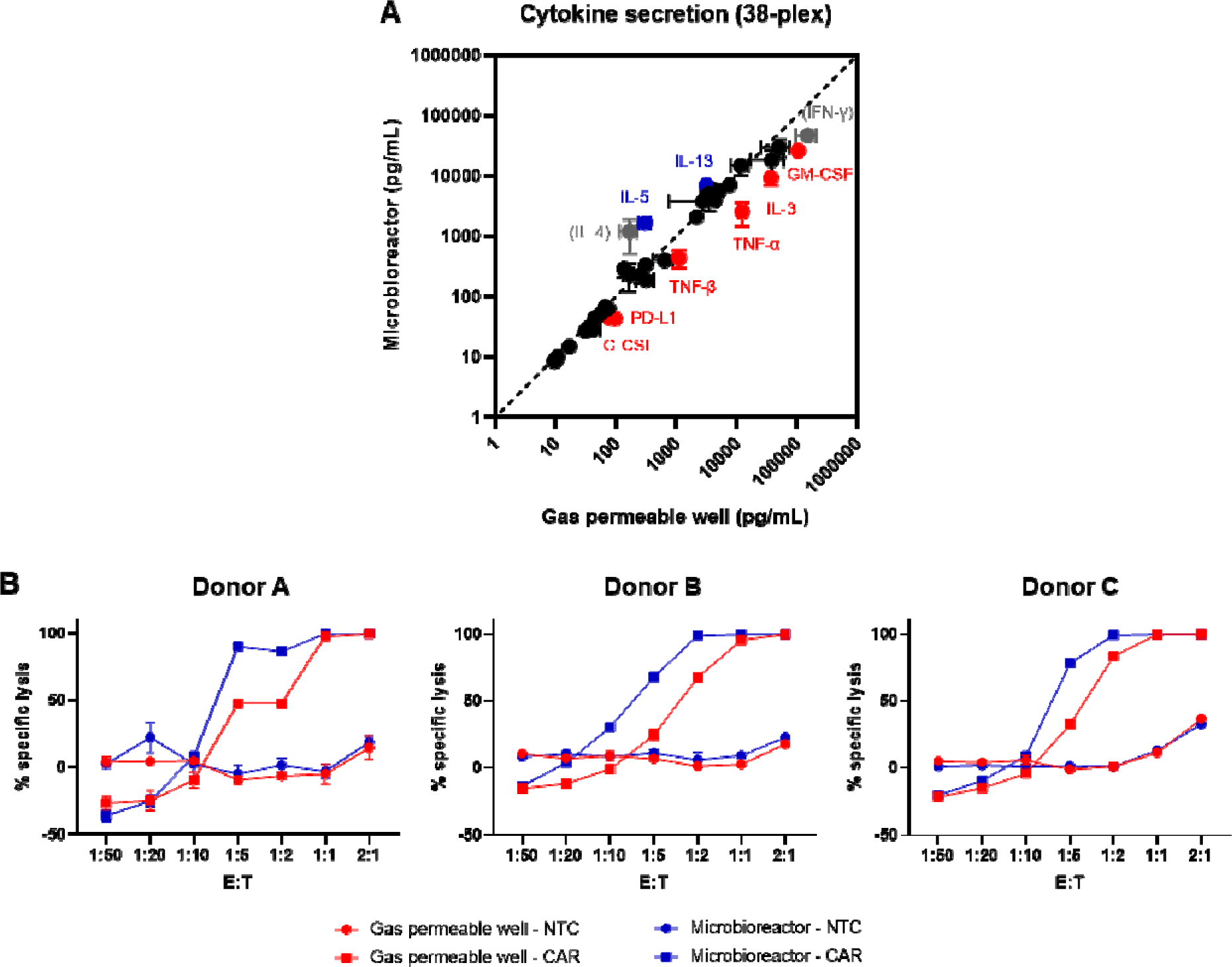
CAR T cells produced on the microbioreactor were highly functional in terms of cytokine release and cytolytic activity against tumor cells. (A) Cytokine secretion following overnight co-culture of T cells and NALM6 cells at a 1:1 CAR+ T cell:NALM6 cell ratio. G-CSF, GM-CSF, IL-3, PD-L1, TNF-L, and TNF-β (red) were secreted at higher levels by T cell from gas-permeable wells, whereas IL-5 and IL-13 (blue) were secreted at higher levels by T cells from the microbioreactor. IFN-γ and IL-4 (grey) were also differentially secreted, albeit not statistically significant. Cell culture supernatant from the co-culture of cells from one technical replicate (one well/cassette) of each donor were used for this assay. Each data point is Mean ± SEM of n=3 donors, each assayed in technical triplicates. Red and blue data points are * P ≤ 0.05 by unpaired t test with Holm-Sidak’s multiple comparisons correction. (B) Percentage specific lysis of NALM6-Luciferase cells following overnight co-culture of T cells and NALM6-Luciferase cells at various CAR+ T cell:NALM6-Luciferase cell ratios. Cryopreserved cells from one technical replicate (one well/cassette) of each donor were thawed for this assay. Percentage specific lysis was determined by luciferase luminescence assay and calculated relative to wells with NALM6 alone (0% lysis). Mean ± SD of n=3 technical replicates.

Anti-CD19 CAR T cells (and non-transduced T cells) were then tested for antigen-specific cytotoxicity. T cells were harvested from the microbioreactor and gas-permeable wells, and cryopreserved at the end of the 12-day process. To evaluate the anti-tumor activity of these cells, we subsequently thawed the cells and co-cultured them with either NALM6 or NALM6-luciferase cells at various effector:target (E:T) cell ratios, following which the number of remaining NALM6 cells were analyzed by flow cytometry cell counting or luciferase luminescence assays, respectively. CAR T cells from all three donors exhibited specific cytolytic activity against tumor cells, when compared with non-transduced T cells (**Figure 4B, Supplementary Figure 7K**). Notably, the cytolytic activity of CAR T cells generated on the microbioreactor were at least comparable with that of CAR T cells generated on the gas-permeable wells (**Figure 4B, Supplementary Figure 7K**). Taken together, these data indicate that CAR T cells produced at high cell densities in a small-volume perfusion-based microbioreactor were equally functional in terms of cytokine secretion and cytotoxic activity compared with CAR T cells produced in gas-permeable well plates.

### Perfusion-based microbioreactor enabled efficient nutrient replenishment and waste removal, and consistent culture environments at high cell densities

The high cell densities achieved in the microbioreactor led us to examine the supply of nutrients and the removal of metabolic waste products, since the accumulation of lactate can inhibit T cell proliferation^43^, and tracking of such attributes could lead to online analytics for the monitoring and control of CAR T cell culture. To track the nutrient and waste levels in the culture medium over time, we sampled cell-free medium daily throughout the 12-day process, and measured metabolite levels offline. Despite starting perfusion at 1 vvd from Day 2 to Day 4, and very low cell numbers at this time, glucose and glutamine levels decreased dramatically in the microbioreactor, and conversely, lactate, glutamate, and ammonia levels increased sharply during this time (**Figure 5**), indicating that the cells exhibited a metabolic switch shortly after activation and transduction, wherein glycolysis and glutaminolysis pathways became highly active. The changes in metabolite levels in the gas-permeable wells were delayed compared with the microbioreactor (**Figure 5**), possibly due in part to the larger static culture volume and greater amount of media supplied from Day 2 to Day 6 of the process (**Supplementary Figure 1D**).

**Figure 5:**
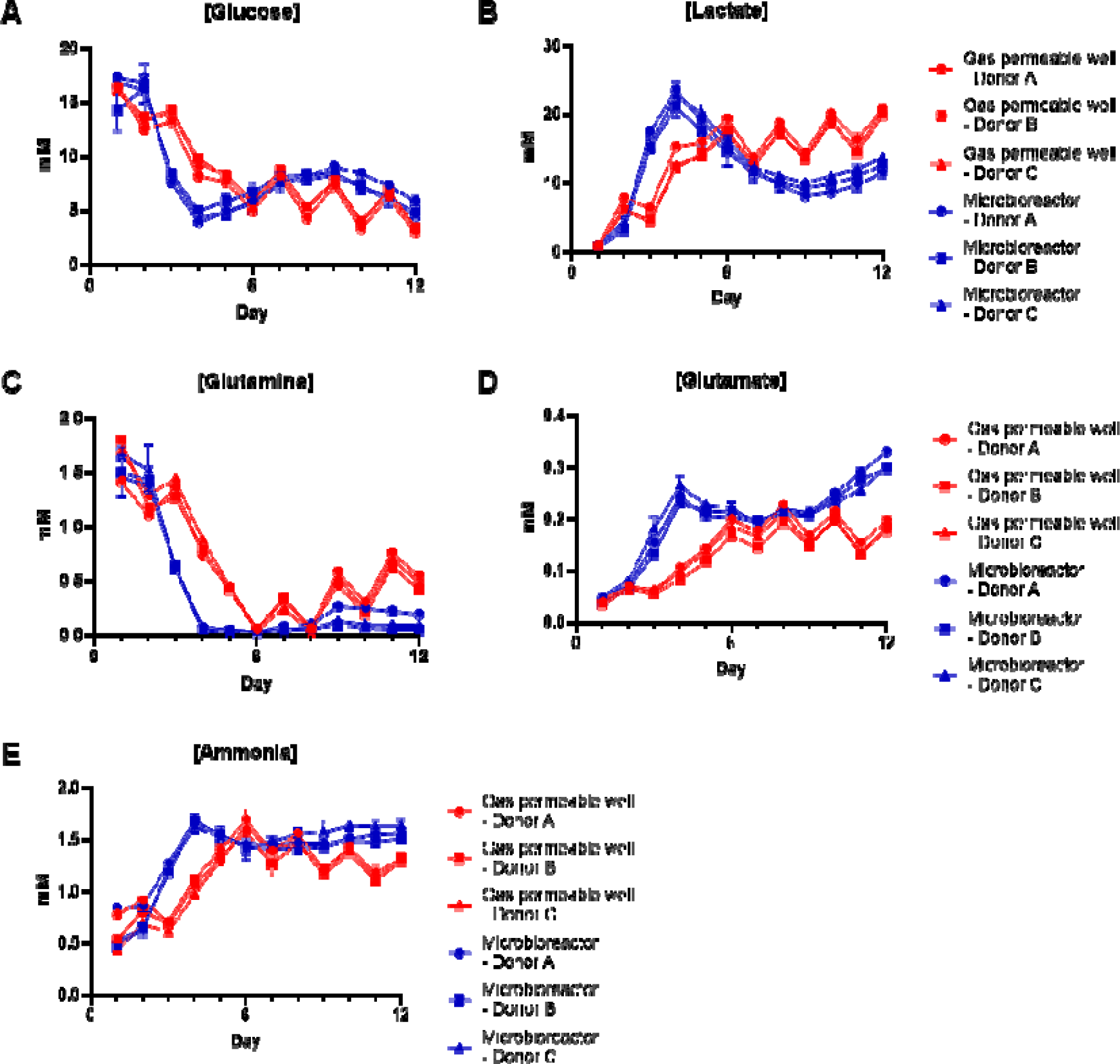
Perfusion-based microbioreactor enabled more efficient nutrient replenishment and waste removal, and more consistent culture environments at high cell densities. To track the nutrient and waste levels in the culture medium over time, cell-free medium from the gas-permeable wells and perfusate from the microbioreactor were sampled daily throughout the 12-day process. Glucose (A), Lactate (B), Glutamine (C), Glutamate (D), and Ammonia (E) concentrations were measured offline using a Cedex Bio Analyzer (Roche CustomBiotech).

In the gas-permeable wells, periodic medium exchanges every other day resulted in fluctuations in metabolite levels from Day 6 to Day 12 when cell numbers are high (**Figure 5**). In contrast, in the microbioreactor, metabolite levels were more consistent due to continuous medium circulation (**Figure 5**). Glucose levels in the microbioreactor were at or above the peak glucose levels in the gas-permeable wells (**Figure 5A**), while lactate levels were lower than that in the gas-permeable wells (**Figure 5B**), indicating more efficient replenishment of glucose and removal of lactate in the microbioreactor, and possibly lower glycolytic activity of the cells in the microbioreactor. In contrast, glutamine levels in the microbioreactor were still lower, and glutamate and ammonia levels were still higher, than those in the gas-permeable wells (**Figure 5C-E**), suggesting that cells in the microbioreactor had higher flux through the glutaminolysis pathway, compared with cells in the gas-permeable wells. Taken together, these data show that the culture environment is highly dynamic, where nutrient availability and waste accumulation may influence the metabolic state of the cells in culture, and vice versa.

### Online process data and offline metabolite measurements enabled modeling of CAR T cell proliferation and metabolic status

There is limited information to date on the cellular metabolic changes over the course of a CAR T cell expansion process^44^. Since the microbioreactor uniquely provides continuous onboard sensor data, such as optical density (OD) as a proxy for total cell density, as well as dissolved oxygen measurements (functionalities that are absent for the gas-permeable well plate), we used these online sensor data together with the offline metabolite measurements to estimate the instantaneous growth rates and metabolic rates of the cells in the microbioreactor (**Figure 6A**). The OD measurements were calibrated by fitting a quadratic equation to the offline cell counts and online OD measurements from all donors and replicates (a global quadratic fit) (**Supplementary Figure 8A**). The imperfection of the fit results in a small uncertainty in the online estimate of cell density. We express this uncertainty by plotting families of curves over the range of calibration parameters (**Supplementary Figure 8B**). These families of curves were applied to the OD data to yield estimates of viable cell counts (VCCs).

**Figure 6:**
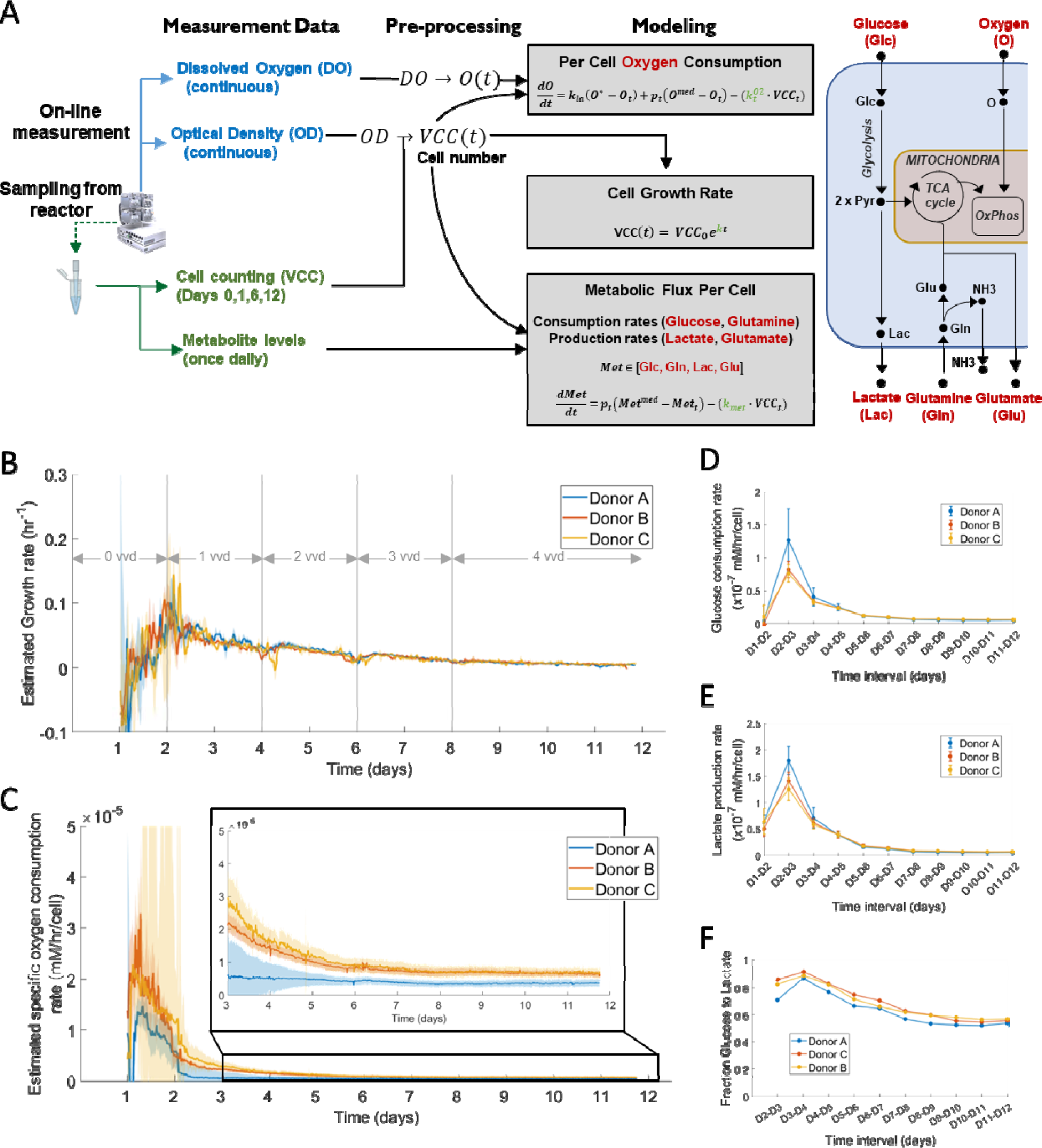
Estimating growth rates and metabolic rates using computational modeling. (A) Schematic showing the flow of input and output data for modeling and the intervening processing steps. The grey boxes in the Modeling column include the equations used to estimate the required rate variables (in green). Further details about the equations can be found in the Methods. The final column shows simplified metabolic pathways that correspond to the modelled metabolites. Cells take in glucose that is processed via glycolysis to yield two molecules of pyruvate. Pyruvate can either be funneled into the Tricarboxylic acid cycle (TCA cycle) for further metabolism, or be converted into lactate and effluxed. The TCA cycle can also be fed via glutamine that loses an amine group to give glutamate and ammonia. Ammonia is typically exported while glutamate can either feed into the TCA cycle (via α–ketoglutarate, not shown) or be effluxed. The TCA cycle leads into oxidative phosphorylation (OxPhos) for oxygen-dependent ATP production. (B-C) Plots of estimated growth and oxygen consumption rates for the three donors over time in the microbioreactor. Median values are plotted as trendlines for each donor. The shaded regions (light blue, pink and yellow) indicate the spread around the median. For each timepoint, the median and the spread around the median (median absolute deviation) were obtained from the three replicates per donor, and from 50 different OD-VCC fit equations per replicate (chosen at random from the family-of-fits for that donor/replicate). (B) shows estimated growth rates (in units of hr^−1^). Grey lines in the plot area correspond to the approximate times when perfusion rate is increased (Day 2 = 1 vvd, Day 4 = 2 vvd, Day 6 = 3 vvd, and Day 8 = 4 vvd). Exact times of perfusion rate changes appear in **Supplementary Table 1**. (C) shows estimated specific oxygen consumption rates (in units of mM O_2_/cell/hr). Inset shows a zoomed in version of the same plot between Days 3 and 12. (D) Plots showing the estimated 24-hour average glucose consumption rates (x10^−7^ mM glucose/cell/hr) for the three donors from Days 1-12. Model-based estimation used the workflow in (A) and 50 different OD-VCC fit equations per replicate (chosen at random from the family-of-fits for that donor/replicate). The x-axes of (D)-(F) show 24-hour time intervals (Day1-Day2, Day2-Day3, etc.). (E) Same as (D) but for lactate production rate (x10^−7^ mM lactate/cell/hr). (F) Plot showing the apparent fraction of glucose funneled towards lactate for the three donors from Days 1-12. This is computed as the model-estimated mean lactate production rate divided by (2*model-estimated mean glucose consumption rate). The factor of two accounts for two lactate molecules being produced from one glucose molecule.

Growth rates (i.e., net rates of proliferation) were computed from the estimated VCCs, and showed the following trends across all three donors: immediately following activation, growth rates increased to a peak of approximately 0.1 hr^−1^, followed by a gradual decline (**Figure 6B**). Then, each time the perfusion rate increased (vvd increased on Days 4, 6, 8) (**Supplementary Figure 8C, Supplementary Table 1**), the growth rate increased transiently, indicating that some cellular proliferation had become limited by nutrient availability or waste accumulation. By Day 12, the estimated growth rates for all donors converged to final values of 0.004-0.005 hr^−1^ (**Figure 6B**). Note that all growth rates are for the total population of cells within a pod, and the growth rates for the underlying phenotypic subsets (Tn/Tscm, Tcm, Tem, Temra) could differ.

Because oxygen is rapidly consumed by high density cultures, the microbioreactor injects oxygen gas in response to a sensor for dissolved oxygen. The data from the oxygen sensor and the oxygen injection rates were analyzed to yield an estimate of the specific (or per cell) oxygen consumption rates over time for each donor (starting from Day 0) (**Figure 6C**). As expected, the oxygen consumption rate peaked immediately after activation, then decreased steadily – providing an independent monitor for the activation protocol. While increases in perfusion rates on Days 4, 6, 8 triggered transient increases in growth rates, the specific oxygen consumption rate did not jump, suggesting that glycolysis fueled the subset of cells that had perfusion-limited expansion. By Day 6, the specific oxygen consumption rates for all donors had reached a plateau with no significant differences until the end of the experiment (Day 12).

To monitor metabolic fluxes in the microbioreactor, we measured metabolites in spent media and constructed a computational model for basic T cell metabolism. The model is a system of differential equations (see Methods) describing the consumption of glucose and glutamine, and the production of lactate and glutamate, per cell per day (starting from Day 2 measurements). To describe uncertainty in calibration for the online cell density, the modeling uses repeated simulations across calibration parameters (families of curves, for each donor and pod). The families of simulations revealed some consistent trends: the consumption of glucose and glutamine increased sharply between Days 2 and 3, then decreased gradually until Day 6, and leveled out until Day 12 (**Figure 6D**, **Supplementary Figure 8D**). Peak rates of glucose consumption were in the range of 0.75-1.3×10^−7^ mM glucose/cell/hr (**Supplementary Table 2**). This peak occurred in the Day2-Day3 interval, which coincided with the peak growth rates (**Supplementary Figure 8G**). Final glucose consumption rates (Day11-Day12) were down to 0.05-0.062 ×10^−7^ mM glucose/cell/hr (**Supplementary Table 2**). Lactate production showed a similar peak between Days 2 and 3 and a gradual decline thereafter (**Figure 6E**). A similar decline was apparent for glutamine consumption and glutamate production (**Supplementary Figure 8D-E**). More importantly, the fraction of glucose shuttled toward lactate production also changed, indicating that the cellular population was more glycolytic on average after activation than at Day 12. At Day 12, only about 60% of the consumed glucose was converted to lactate and the rest went to other metabolic or proliferative processes (such as oxidative phosphorylation) (**Figure 6F**). In contrast, the fraction of glutamine funneled towards glutamate increased over time, suggesting that cells were performing increasing amounts of glutaminolysis over the course of the 12-day period (**Supplementary Figure 8F**). There is a good quality of fit between model-estimated metabolite concentrations and measured metabolite concentrations (**Supplementary Figure 9, 10**). Taken together, these data show that the metabolic profile of the cells changes over time in culture, and could be a useful attribute to monitor and control in a CAR T cell production process.

## Discussion

Current bottlenecks in autologous cell therapy manufacturing, such as lengthy processes, large equipment footprints, low production throughputs, need for centralized cleanroom facilities, and high cost of goods, would benefit from the tenets of process intensification. When applied to other biomanufacturing areas, such as in monoclonal antibody production, perfusion-based continuous manufacturing at high cell densities have maximized productivity and cost-efficiency while minimizing resource usage and manufacturing costs^45, 46^. Here, we demonstrate the high-density production of functional anti-CD19 CAR T cells in a perfusion-capable 2 mL microfluidic bioreactor, with better cell expansion compared with a static fed-batch gas-permeable well plate.

With improved medium exchange and gas transfer through the continuous perfusion of medium and injection of O_2_ into the headspace to maintain dissolved O_2_ levels, respectively, T cell expansion was enhanced in the microbioreactor, particularly from Day 6 to Day 12 of the process, when glucose levels could be limiting and lactate levels could be inhibitory^43^ in the gas-permeable wells. Other forms of fed-batch culture systems^20, 44, 47^ also suffer from relatively low T cell expansion rates. With high T cell expansion rates, similar total T cell numbers could be attained with a shorter culture period in the microbioreactor (7-8 days) compared with the gas-permeable wells (12 days), thus representing a potential shortening of production runs by 30-40%. This could substantially reduce the turnaround times and increase the number of CAR T cell production slots, thus raising manufacturing capacity. This is also in line with the industry’s shift towards shorter CAR T cell culture periods, which have been shown to result in cell therapy products with less-differentiated phenotypes, higher functionality and longer *in vivo* persistence^3, 17, 48^. The high T cell numbers in the small-volume microbioreactor was possible due to its ability to support viable cell densities of close to 200 million cells/mL, which is at least an order of magnitude greater than what is achievable in the gas-permeable wells, as well as in other culture systems^20, 22, 23, 44^. Currently, only hollow-fiber bioreactors, such as the Quantum Cell Expansion System (Terumo BCT)^24^, are able to achieve similarly high viable cell densities. However, in contrast to the microbioreactor, the Quantum is a comparatively large volume bioreactor, with intracapillary and extracapillary circulation volumes of at least 100 mL and 300 mL, respectively, that requires a high total medium usage of more than 10 L per run^31^. Furthermore, as cells are inoculated within the intracapillary circulation, viral vector transduction, in-process cell sampling, and final cell harvest can be complicated to perform^22^. In contrast, these manipulations are straightforward to perform on the microbioreactor via the inoculation, sampling, and sterile waste ports, respectively.

Conversely, use of the Breez alleviates many of the drawbacks of large-scale bioreactors, such as large equipment footprints, low production throughputs, and the need for centralized cleanroom facilities. For example, rocking bag bioreactors, and the CliniMACS Prodigy (Miltenyi Biotec), have minimum working volumes of 100-200 mL and equipment footprints of 0.3-0.4 m^2^. Furthermore, they are able to process only a single sample at a given time. In contrast, the scaled-down microbioreactor used in this study is able to run up to four independent cultures, either simultaneously or asynchronously, in a small form factor of 0.2-0.3 m^2^, hence allowing flexible and highly-parallelized CAR T cell cultures. The volume reduction required low seeding cell numbers, which in turn resulted in low LVV and medium usage compared with other bioreactors. The reduced LVV usage and improved medium efficiency per unit number of CAR T cells produced could be advantageous in the context of limited supplies and/or high costs of GMP-grade viral vectors and reagents, such as medium, serum, and cytokines used for CAR T cell manufacturing^49–51^. Moreover, the automated closed-system microbioreactor can be operated on the bench top with lower manpower and no cleanroom requirements, thereby reducing labor and cleanroom costs, which are significant fixed costs in CAR T cell production^11^, and thus potentially enabling decentralized, point-of-care manufacturing of CAR T cells at the bedside. The process intensification enabled by the advanced integrated microfluidics of perfusion microbioreactors could substantially improve space-time yields and decrease the cost of goods of adoptive cell therapies.

Apart from being potential future manufacturing platforms, scaled-down microbioreactors could also be valuable process development tools. There is a general lack of process understanding in cell therapy manufacturing^17, 52, 53^, partly due to the lack of suitable manufacturing technologies at the appropriate scale. This microbioreactor, with four pods per system, and on-board environmental measurements that allow greater process control and predictability, could facilitate future high-throughput process characterization under controlled conditions. Probing a larger process design space, such as through Design of Experiments (DoE) or systems modeling methodologies according to Quality by Design (QbD) principles, could lead to a better understanding of how multivariate process parameters and biological factors intersect and impact upon CAR T cell quality attributes^13, 52^. This would aid in process optimization studies, such as for different cell therapy products, which could have unique metabolic characteristics^54^, or for different donors, for which considerable biological heterogeneity is well recognized^55, 56^.

Examples of such donor variability were evident in our study. Donor B exhibited lower proportions of Tn/Tscm cells and higher proportions of Tcm and Tem cells at the end of the process, independent of the culture vessel, compared with Donors A and C. On the other hand, Donor C showed markedly higher proportions of CD57+ cells in the microbioreactor but not in the gas-permeable wells. Overall, microbioreactor cultures resulted in slightly higher proportions of more-differentiated Tem and Temra cells (and lower proportions of less-differentiated Tn/Tscm and Tcm cells) at the end of the process, compared with gas-permeable well plate cultures. Part of this might be due to the higher rates of cell expansion in the microbioreactor compared with the gas-permeable wells, particularly from Day 6 to Day 12 of the process, since these phenotypic differences were less apparent on Day 6, when fold expansion and viable cell counts were just slightly higher in the microbioreactor compared with the gas-permeable wells. However, other factors, such as starting cellular phenotype^57^, level of activation^56^, and cytokine signaling^17^, can also influence the differentiation status of *ex vivo* T cell cultures. Optimizing process parameters, including the timing, identity, and dosages of growth factors, media, and activation agents used in culture, could be beneficial for further improving cell yields and phenotypes in perfusion microbioreactors.

In this study, we demonstrated a proof-of-concept using online optical density (OD) and dissolved O_2_ (DO) measurements, as well as offline metabolite measurements, to derive cell densities, per-cell oxygen consumption rates as an indicator of oxidative phosphorylation rates, as well as per-cell glucose/glutamine consumption rates and lactate/glutamate production rates as indicators of glycolysis/glutaminolysis rates, respectively, to monitor shifts in T cell metabolism during *ex vivo* cultures. While differences in glycolysis/glutaminolysis rates and T cell differentiation phenotypes between donors were relatively subtle in our study, other studies have suggested that inhibition of glycolysis/glutaminolysis can uncouple T cell expansion from T cell differentiation, thus favoring the formation of less-differentiated naïve and central memory T cells, which are associated with longer *in vivo* persistence^58–60^. In this regard, future integration of online metabolite sensors, such as using Raman spectroscopy^61^, could enable in-process monitoring of CAR T cell metabolic fitness. An enhanced process knowledge, such as of the metabolic changes during *ex vivo* culture from the data gathered in this study, could form the basis of future feedback-driven processes, such as adjusting perfusion rates based on metabolic state instead of a fixed protocol, to provide consistent culture conditions for CAR T cell growth, or even to drive the cells toward a more desirable phenotype, while minimizing the volume of medium used^62, 63^. In summary, the advanced bioprocess controls enabled by CAR T cell culture-on-a-chip could pave the way for future adaptive manufacturing processes^64^ that can mitigate starting material variability and result in cell therapies with improved consistency and efficacy for greater patient benefit.

### Methods Cell lines

Lenti-X 293T Cell Line was purchased from Takara Bio Inc., and maintained in HyClone Dulbecco’s Modified Eagle Medium (DMEM) with high glucose (Cytiva) supplemented with 10% Fetal Bovine Serum (Thermo Fisher Scientific) and 1% Penicillin-Streptomycin (Thermo Fisher Scientific). Jurkat, Clone E6-1 (TIB-152) and NALM6, clone G5 (CRL-3273) cell lines were purchased from ATCC, and maintained in HyClone RPMI 1640 medium (Cytiva) supplemented with 10% Fetal Bovine Serum (Thermo Fisher Scientific) and 1% Penicillin-Streptomycin (Thermo Fisher Scientific).

### Plasmids

pMDLg/pRRE, pRSV-Rev, and pMD2.G packaging plasmids were gifts from Didier Trono (Addgene plasmid # 12251, 12253, and 12259)^65^. A second-generation anti-CD19 CAR-IRES-EGFP transfer plasmid, encoding a codon-optimized anti-CD19 CAR comprised of Myc-epitope-tagged FMC63 single-chain variable fragments, IgG4 hinge, CD28 transmembrane domain, and human 4-1BB and CD3zeta intracellular signaling domains, was generated as described previously^66^. Research grade plasmid DNA was prepared by GenScript Biotech.

### Lentiviral vector production and titration

Lentiviral vectors were generated by first transfecting the Lenti-X 293T Cell Line with the packaging plasmids and transfer plasmid complexed with Lipofectamine 3000 Transfection Reagent (Thermo Fisher Scientific). Medium was changed 4-6 h after transfection, and lentiviral supernatants were collected 48-96 h after transfection. The supernatant was then centrifuged for 5 min at 300g, 4°C, and filtered through a 0.45-μm low-protein-binding filter. The clarified supernatant was then ultracentrifuged for 1.5 h at 75,000g, 4°C. The lentiviral pellet was then resuspended in Opti-MEM Reduced Serum Medium (Thermo Fisher Scientific) at 4°C, and stored in aliquots at −80°C. The concentrated lentiviral vectors were titrated on Jurkat cells. The percentages of GFP+ Jurkat cells were measured by flow cytometry two days post-transduction, and lentiviral titers were calculated and expressed in TU/mL.

### Preparation and operation of microbioreactor

Prior to starting a production run, medium preparation, system set up, and automated priming and calibration of the microfluidic cassettes were performed. This process primes the fluid lines, calibrates the optical density, dissolved O_2_, and pH sensors, and calculates the volumetric mass transfer coefficient (kLa) of the cassettes. Cell inoculation was performed by connecting a syringe containing the cell-bead mixture to the inoculation port in a BSC. The cassette was then loaded onto the pod, and the cell-bead mixture pulled into the growth chamber by vacuum inoculation. For transduction, 1 mL of cell-free medium was removed through the perfusion output, and then 1 mL of fresh medium containing diluted LVV was pulled into the growth chamber by vacuum inoculation, through a syringe connected to the inoculation port, analogous to the process for cell inoculation. The growth chamber was continuously mixed for 2 h after the addition of LVV, and then subsequently reverted to intermittent mixing for the remainder of the process. The microbioreactor cultures were controlled at the following set-points: temperature of 37°C, minimum CO_2_ levels of 5%, minimum dissolved O_2_ levels of 80% air saturation, and a pH of 7.40 ± 0.05 (to reflect the physiological definition of acidosis and alkalosis). Dissolved O_2_ control was implemented such that when dissolved O_2_ levels fall below the setpoint of 80%, O_2_ was injected into the headspace of the cassette to maintain dissolved O_2_ levels at a minimum of 80%. pH control was implemented such that when pH is above 7.45, more CO_2_ (above the minimum of 5%) would be injected into the headspace of the cassette to lower the pH, and when pH is below 7.35, a basic carbonate/bicarbonate solution (Bioreactor pH Adjustment Solution, Merck Sigma-Aldrich) would be injected into the growth chamber to raise the pH. For cell harvest, the sterile waste bottle was rinsed with phosphate-buffered saline (PBS) and emptied in a BSC. The cassette was then loaded onto the pod, and the contents of the growth chamber ejected into the waste bottle, followed by removal of the cell suspension from the waste bottle using a syringe in a BSC. Apart from cell or viral vector inoculation and cell harvest in a BSC, the rest of the process occurs on the system placed on the bench top in an academic tissue culture room facility. Cell samples were taken by sampling 50 uL through the sampling port into a 1.5-mL tube. Cell-free medium samples were taken by removal of ∼ 200 uL perfusate (accumulated over ∼ 2.4 h for 1 vvd, 1.2 h for 2 vvd, 48 min for 3 vvd, and 36 min for 4 vvd) from the perfusion output using a syringe.

### Human T cell isolation, activation, transduction, and expansion

Frozen human peripheral blood mononuclear cells from healthy donors were purchased from STEMCELL Technologies. T cells were isolated using EasySep Human T Cell Isolation Kit (STEMCELL Technologies), and resuspended in AIM V Medium (Thermo Fisher Scientific) supplemented with 2% human male AB serum (Merck Sigma-Aldrich) and 100 IU/mL recombinant human IL-2 (Miltenyi Biotec). For activation, purified T cells were mixed with DynaBeads Human T-Expander CD3/CD28 (Thermo Fisher Scientific) at a 1:1 cell:bead ratio in a 25 mL tube, and 2 mL of cell-bead mixture was seeded into each gas-permeable well of a G-Rex 24-well plate (Wilson Wolf) and each microbioreactor cassette (Erbi Biosystems, MilliporeSigma). One day after activation, 1 mL of cell-free medium was removed, and then 1 mL of fresh medium containing diluted LVV was added to the wells or cassettes at a MOI of 5. One day after transduction, 6 mL of fresh medium was added to achieve the final culture volume of 8 mL for the gas-permeable wells, and perfusion was started at 1 vvd for the microbioreactor. For the gas-permeable wells, 6 mL medium was exchanged every other day, with a constant culture volume of 8 mL per well. For the microbioreactor, the perfusion flow rate was increased by 1 vvd every other day until a maximum of 4 vvd, with a constant culture volume of 2 mL. T cells were expanded over 10 days, for a total process duration of 12 days.

### Cell count and viability

Cell counts and viabilities were measured using Acridine Orange (AO) and Propidium Iodide (PI) and/or Trypan Blue (TB) staining on a CellDrop Automated Cell Counter (DeNovix).

### Cell-free medium metabolite analysis

Metabolites in cell-free medium, including glucose, lactate, glutamine, glutamate, and ammonia, were measured on a Cedex Bio Analyzer (Roche CustomBiotech).

### Flow cytometry

For T cell purity analysis, the panel used include: LIVE/DEAD Fixable Violet (Thermo Fisher Scientific) and PE/Cy7 anti-CD3 (clone HIT3a) (Biolegend). For T cell activation analysis, the panel used include: LIVE/DEAD Fixable Violet (Thermo Fisher Scientific), BV510 anti-CD4 (clone OKT4), BV650 anti-CD8a (clone RPA-T8), PE/Cy7 anti-CD25 (clone BC96), APC anti-CD69 (clone FN50) (Biolegend), and R718 anti-LDL-R (clone C7) (BD Biosciences). For T cell surface phenotyping, the panel used include: LIVE/DEAD Fixable Violet (Thermo Fisher Scientific), BV510 anti-CD4 (clone OKT4), BV605 anti-CD45RA (clone HI100), BV650 anti-CD8a (clone RPA-T8), PerCP/Cy5.5 anti-LAG-3 (clone 11C3C65), PE anti-CCR7 (clone G043H7), PE/Dazzle 594 anti-CD127 (clone A019D5), PE/Cy7 anti-CD57 (clone HNK-1), AF700 anti-PD-1 (clone EH12.2H7), and APC/Fire 750 anti-TIM-3 (clone F38-2E2) (Biolegend). Cells were washed with PBS (Thermo Fisher Scientific), then stained by incubating with LIVE/DEAD Fixable Violet Stain for 20 min at room temperature. Cells were subsequently washed with FACS Buffer [PBS supplemented with 2% Fetal Bovine Serum (Thermo Fisher Scientific) and 0.1% sodium azide (Merck Sigma-Aldrich)], then stained by incubating with antibodies for 20 min at room temperature. Cells were subsequently washed with FACS Buffer again before acquisition on a CytoFLEX S flow cytometer (Beckman Coulter).

### Cytokine secretion analysis

Fresh end-of-production T cells were co-cultured overnight with NALM6 cells at an effector:target (E:T) ratio (CAR+ T cell:NALM6 cell) of 1:1 in AIM V Medium (Thermo Fisher Scientific) supplemented with 2% human male AB serum (Merck Sigma-Aldrich) and 100 IU/mL recombinant human IL-2 (Miltenyi Biotec). Cell culture supernatants were harvested and stored at −80L. Cytokine and chemokine secretion profiles were determined using the Human XL Cytokine Luminex Performance Assay 46-plex Fixed Panel (R&D Systems) and measured on a Luminex FLEXMAP 3D system (Thermo Fisher Scientific). Because Granzyme B and IFN-γ were above the range of the standard curve, these two cytokines were re-analyzed using the Human Granzyme B and IFN-γ ProcartaPlex Kit (Thermo Fisher Scientific) and measured on a Luminex FLEXMAP 3D system (Thermo Fisher Scientific). Out of 46 cytokines that were measured, 38 cytokines had samples that were within range of the standard curve, of which 8 cytokines were significantly differentially secreted – 6 cytokines (G-CSF, GM-CSF, IL-3, PD-L1, TNF-α, and TNF-β) were secreted at significantly higher levels by gas-permeable well CAR T cells when stimulated with NALM6 cells, whereas 2 cytokines (IL-5 and IL-13) were secreted at significantly higher levels by microbioreactor CAR T cells when stimulated with NALM6 cells.

### Cytotoxicity assay

Cryopreserved end-of-production T cells were thawed and co-cultured overnight with either NALM6 cells pre-stained with CellTrace Violet (Thermo Fisher Scientific) for 20 min at 37°C or NALM6-Luciferase cells at various effector:target (E:T) ratios (CAR+ T cell:NALM6 cell) in RPMI 1640 medium (Cytiva) supplemented with 10% Fetal Bovine Serum (Thermo Fisher Scientific) and 1% Penicillin-Streptomycin (Thermo Fisher Scientific). For flow cytometry-based cytotoxicity assay with NALM6 cells, co-cultured cells were collected and washed with FACS Buffer, then stained with PE/Cy7 anti-CD3 (clone HIT3a) (Biolegend) for 20 min at room temperature. Cells were subsequently washed with FACS Buffer again, and SYTOX AADvanced Dead Cell Stain (Thermo Fisher Scientific) and CountBright Plus Absolute Counting Beads (Thermo Fisher Scientific) were added, before acquisition on a CytoFLEX S flow cytometer (Beckman Coulter). For luciferase luminescence-based cytotoxicity assay with NALM6-Luciferase cells, co-cultured cells were transferred to a white 96-well opaque-bottom plate. Bright-Glo Luciferase Assay reagent (Promega) was added to the wells, and luminescence signals were measured using Infinite M200 Pro plate reader (Tecan).

### Metabolic Modeling

#### OD offset correction

After gathering the optical density (OD) values recorded by the microbioreactor for each donor and each replicate, we observed that a few of the runs had many negative OD values at early timepoints (prior to Day 4). This may be due to a high baseline for cell-free media due to bubbles or lamination effects in the OD path during initial cassette calibration. Hence, we performed an OD correction by adding an offset as follows. For every donor/replicate, the offset value was chosen to be the smallest value (>=0) whose addition to the OD would ensure that 5% or less of corrected OD values between Days 0-4 were negative. These offset-corrected OD values were then used for all subsequent modeling and computation.

#### Estimating cell numbers from optical density

We gathered the experimental measurements of viable cell counts (VCCs) from infrequent sampling of each culture, and compared them against the optical density (OD) values for the same timepoint when the cells were sampled from the reactor. We used these paired values of OD and VCC (from all donors and replicates) to create a pooled dataset for converting OD to VCC. We fit the relationship to a quadratic curve:

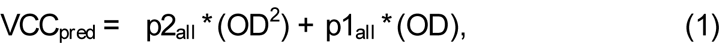

where *p2_all_* is the quadratic coefficient and *p1_all_*is the linear coefficient of the generalized OD-VCC fit. The parameters *p2_all_*and *p1_all_* were fit to the pooled dataset using the MATLAB fit function, and we call this the generalized OD-VCC fit. To further tailor this fit for each donor and replicate (i.e., for each hardware cassette), we fixed the quadratic coefficient at the value of *p2_all_*, and re-fit a new linear coefficient for each donor/replicate using the MATLAB fit function. We call this the linear-customized fit (green curve in **Supplementary Figure 8B**).

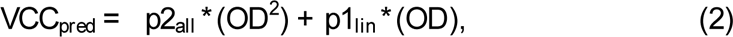

where *p1_lin_* is the newly fit linear coefficient for each donor and replicate. In parallel, we also obtained a completely fresh fit for each donor and replicate, that was performed only using the cell counts and OD values for the specific donor and replicate. We call this the fully-customized fit.

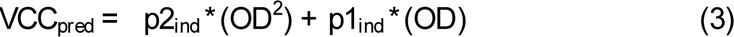

Using the linear-customized fit and the fully-customized fit, we generated a family of fitted curves for each donor and replicate as follows: we generated a new quadratic term by sampling uniformly at random between *p2_all_*and *p2_ind_* while generating a new linear term by sampling uniformly between *p1_lin_* and *p1_ind_*. This was repeated 1000 times. We obtained our final family-of-fits for each donor/replicate by choosing the subset of fit equations (out of the 1000 fits) whose predictions of the VCC fell between the 25^th^ and 75^th^ percentile of the 1000 predictions (grey curves in **Supplementary Figure 8B**). Estimates of the VCC were utilized for subsequent analysis of the growth rates and metabolic rates as described below.

#### Computing instantaneous growth rates

First we processed the OD data for use in inferring cell numbers. We took the microbioreactor OD data (measured roughly once per minute) and smoothed it by averaging over a sliding window of size 21 minutes (10 on each side of an OD value). Because the sampling periods are not exactly 1 minute in each pod, we obtained synchronized readouts of all pods and all runs by interpolating the smoothed averages to obtain OD values at intervals of exactly 0.1 hrs (6 minutes), starting at 24.00 hrs. Each set of OD values was converted to a family of VCC curves, using the method above. Finally, we computed instantaneous growth rates for each donor/replicate as follows. Growth rates were computed between points separated by a one-hour interval, using the difference equation (5) obtained from the simple exponential cell growth equation (4).

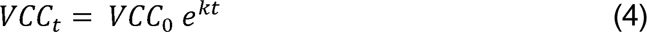

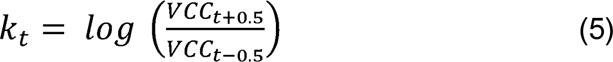

where *k_t_* is the estimated growth rate at time *t,* and *VCC_t+0.5_* and *VCC_t-0.5_* are the estimated VCCs at timepoints *t*-0.5 hrs and *t*+0.5 hrs respectively. We performed a final smoothing of the estimated growth rates using a sliding window of 40 datapoints (4 hr interval). This process was repeated for 50 randomly chosen fit equations from the family-of-fits for that donor/replicate, resulting in a set of 50 growth rate estimates for each timepoint, each donor, and each replicate.

#### Computing specific oxygen consumption rates

To compute specific oxygen consumption rates (oxygen consumption rate per cell), we followed a process similar to the growth rate computation. For each donor/replicate, OD was smoothened using a sliding window of size 21, followed by interpolation to obtain OD values for timepoints separated by an interval of 0.1 hrs (6 minutes) starting from the 24 hr timepoint. Specific oxygen consumption rates were computed from the difference equation (7) based on the ODE in (6).

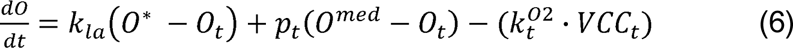

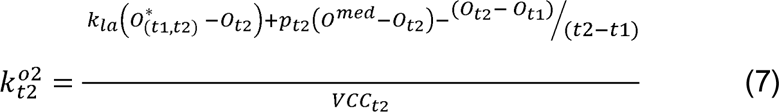

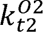 is the specific oxygen consumption rate at time *t2*; k_la_ is the volumetric mass transfer coefficient for oxygen for the microbioreactor cassette (obtained during cassette calibration); O_t1_ and O_t2_ are the dissolved oxygen concentrations at timepoints *t1* and *t2* (*t2*>*t1*); p_t2_ is the perfusion rate at time *t2*; O^med^ is the concentration of dissolved oxygen in fresh media and VCC_t2_ is the estimated VCC at time 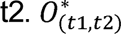 is the saturating oxygen concentration in the time interval between *t1* and *t2*, and is computed using the fraction of time pure O_2_ vs air is supplied via the oxygen controller, in the time interval from t1 to t2. Similar to the growth rate computation, we repeated the specific oxygen consumption rate computation with 50 randomly chosen fit equations from the family-of-fits for that donor/replicate. This workflow yields a set of 50 estimates of specific oxygen consumption rate for each timepoint, each donor, and each replicate.

#### Computing daily metabolite consumption/production rates

Daily metabolite consumption/production rates were computed using the OD values from the microbioreactor and offline measurements of metabolites (once per day) in perfusate media from the microbioreactor. In particular, we computed the average daily consumption rates of glucose and glutamine per cell, and average daily production rates of lactate and glutamate per cell. For each donor and each replicate, we computed a 24hr average for the following four metabolic rates: glucose consumption, glutamine consumption, lactate production, and glutamate production (all using units of mM per cell per hour). The process of estimating metabolic rates uses ordinary differential equations (ODEs) of the following form

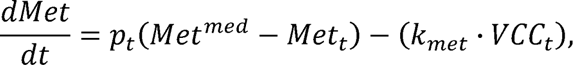

where 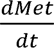 represents the rate of change of the metabolite concentration over time, p_t_ is the perfusion rate at time t, Met^med^ is a constant indicating the concentration of the metabolite in fresh media, Met_t_ is the (unknown) concentration of the metabolite in the microbioreactor at time (t), VCC_t_ is the estimated VCC at time *t*, and k_met_ is the unknown rate of metabolite consumption/production. The metabolite concentrations over time in the reactor (Met_t_) can be computed easily with an ODE solver if we know the values of p_t_, Met^med^, k_met_, VCC_t_, and the initial value of Met_t_. We do not know k_met_, but we can estimate it using repeated guesses and repeated attempts of ODE simulation, based on the expectation that Met_t_ should resemble the offline metabolite measurements in spent media. In other words, we optimized k_met_ while minimizing an objective function that forced the simulated value of Met to match the offline measurements of Met (at the timepoints when spent media was collected). The objective function used a sum of squares penalty as follows.

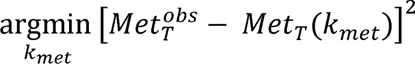

Where *T* represents the times at which spent media was collected for offline measurements. The optimization was performed independently for 11 time-intervals between successive experimental measurements of the metabolite concentrations (Day1-Day2, Day2-Day3,…., Day11-Day12). The optimization method was simulated annealing using the MATLAB command *simulannealbnd*, and the ODE solver was MATLAB *ode15s.* Note that the ODE simulations are actually families of simulations, because we estimate VCC_t_ by taking 50 randomly chosen fit equations from the family-of-fits for that donor and replicate. To avoid local minima, we repeated the entire optimization process 5 times, each starting with a different random seed for the initialization of k_met_ in the optimizer. This workflow yields an optimized value of k_met_ for each day, metabolite, donor, and replicate, meaning that the metabolic rates of the system are represented by step functions, with a discontinuity every 24hrs. Figures were plotted by combining all replicates for each donor, and the error bars include the 50 fit equations for each replicate. In other words, 150 points go into the error bar for each donor with 3 replicates.

### Statistical analysis

Statistical analyses were performed using the Prism 9 (GraphPad) software. For statistical comparisons, significance was determined using two-tailed unpaired parametric t-tests. Adjusted P values < 0.05 after multiple comparison correction, where required, were considered statistically significant. The statistical test used for each experiment is noted in the relevant figure legend.

## Data availability

All data generated or analyzed during the study are included within the paper and its supplementary information.

## Code availability

The code used for computational modeling can be accessed at https://github.com/nsuhasj/perfReactorCarT.

## Acknowledgements

This research is supported by the National Research Foundation, Prime Minister’s Office, Singapore under its Campus for Research Excellence and Technological Enterprise (CREATE) programme, through the Singapore-MIT Alliance for Research and Technology (SMART) Critical Analytics for Manufacturing Personalized-Medicine (CAMP) Inter-Disciplinary Research Group. The authors would like to thank Ms. Veonice Au of the Institute of Molecular and Cell Biology (IMCB), Agency for Science Technology and Research (A*STAR) for assistance with Luminex multiplex assays, and Dr. Kevin S. Lee of Erbi Biosystems, part of MilliporeSigma, for technical assistance with the microbioreactor.

## Author contributions

W.-X.S., R.J.R., L.T.-K., and M.E.B. conceived of the project. W.-X.S., D.B.L.T., F.K., D.S., and K.-W.C conducted experiments. W.-X.S., N.S.J., D.B.L.T., and K.-W.C. analyzed data. Y.H.L., R.J.R., L.T.-K., and M.E.B. supervised the work. W.-X.S., N.S.J., L.T.-K., and M.E.B. wrote the manuscript. All authors edited the manuscript.

