## Supplemental Information for "High-density microbioreactor process designed for automated point-of-care manufacturing of CAR T cells"

[1] Critical Analytics for Manufacturing Personalized-Medicine (CAMP), Singapore-MIT Alliance for Research and Technology (SMART), Singapore.
[2] Cancer and Stem Cell Biology, Duke-NUS Medical School, Singapore.
[3] Department of Electrical Engineering and Computer Science, Massachusetts Institute of Technology, Cambridge, MA, USA.
[4] Department of Biological Engineering, Massachusetts Institute of Technology, Cambridge, MA, USA.
[5] Koch Institute for Integrative Cancer Research, Cambridge, MA, USA.
[6] Ragon Institute of MIT, MGH and Harvard, Cambridge, MA, USA.

* Corresponding authors:
Michael E. Birnbaum:
Lisa Tucker-Kellogg:
Rajeev J. Ram:

Short title:
CAR T cell culture-on-a-chip

Keywords:
High-density; CAR T cells; Perfusion; Microfluidic; Bioreactor

### Supplementary Information

10 Supplementary Figures
2 Supplementary Tables

A

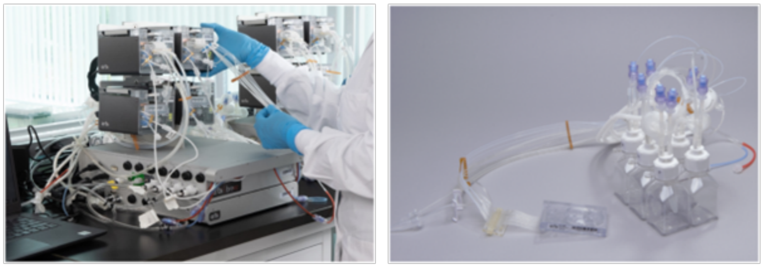

© 2023 Merck KGaA, Darmstadt, Germany and/or its affiliates. All rights reserved

B

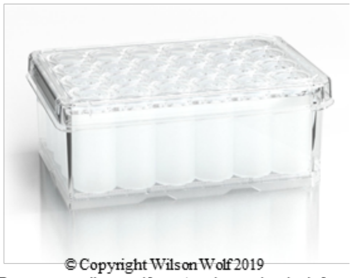

© Copyright Wilson Wolf 2019
From: www.wilsonwolf.com/product-and-order-info

C

| Step / Parameter | G-Rex® 24 well plate | Breez™ microbioreactor |
|---|---|---|
| **Activation** | 2 million cells + 2 million Dynabeads in 2 mL medium | |
| **Transduction** | Remove 1 mL medium, then add 1 mL medium containing LVV at MOI 5 | |
| **Culture volume** | 8 mL | 2 mL |
| **Medium exchange** | 6 mL every other day | Continuous perfusion at 1 vvd to 4 vvd |
| **Total medium volume used** | 33 mL | 59 mL |
| **Temperature control** | 37°C incubator | 37°C |
| **$CO_2$ control** | 5% incubator | 5% (minimum) |
| **$O_2$ control** | No control | 80% air saturation (minimum) |
| **pH control** | No control | 7.40 ± 0.05 |

D

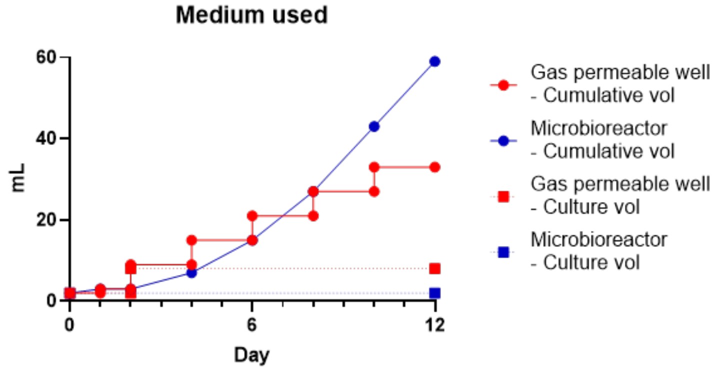


E

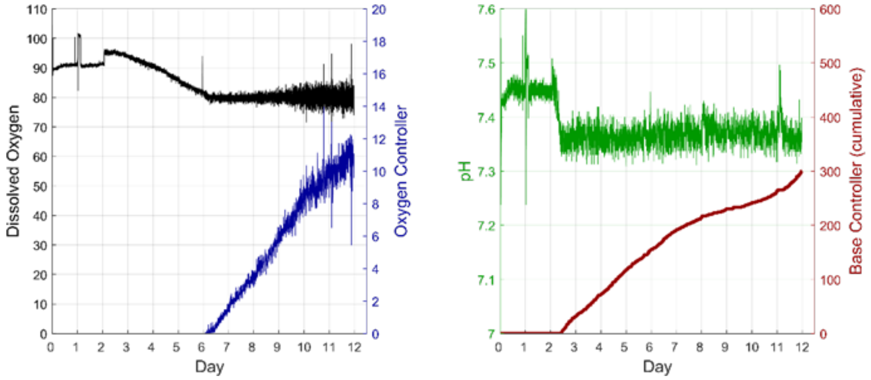


**Supplementary Figure 1: Details of CAR T cell production process on the microbioreactor and gas-permeable well plate.** (A) Images of the microbioreactor system, comprised of a base station controller and a $CO_2$ controller supporting up to four "pods" (left), and of the sterile, singleuse consumable, consisting of a microfluidic chip linked to a bottle rack assembly (right). (B) Image of the gas-permeable well plate. (C) Details of activation, transduction, and expansion culture conditions on the microbioreactor and gas-permeable wells. (D) Culture volumes (dotted lines) and total volumes of medium used (solid lines) for the microbioreactor and gas-permeable wells. (E) The microbioreactor dissolved $O_2$ setpoint was 80%. When dissolved $O_2$ levels fall below 80%, $O_2$ would be injected into the headspace of the cassette to maintain dissolved $O_2$ levels at a minimum of 80%. The microbioreactor pH setpoint was 7.40 ± 0.05. When pH is above 7.45, more $CO_2$ (above the minimum of 5%) would be injected into the headspace of the cassette to lower the pH, and when pH is below 7.35, a basic carbonate/bicarbonate solution would be injected into the growth chamber to raise the pH. Dissolved $O_2$ and pH control were activated 2 h after transduction.

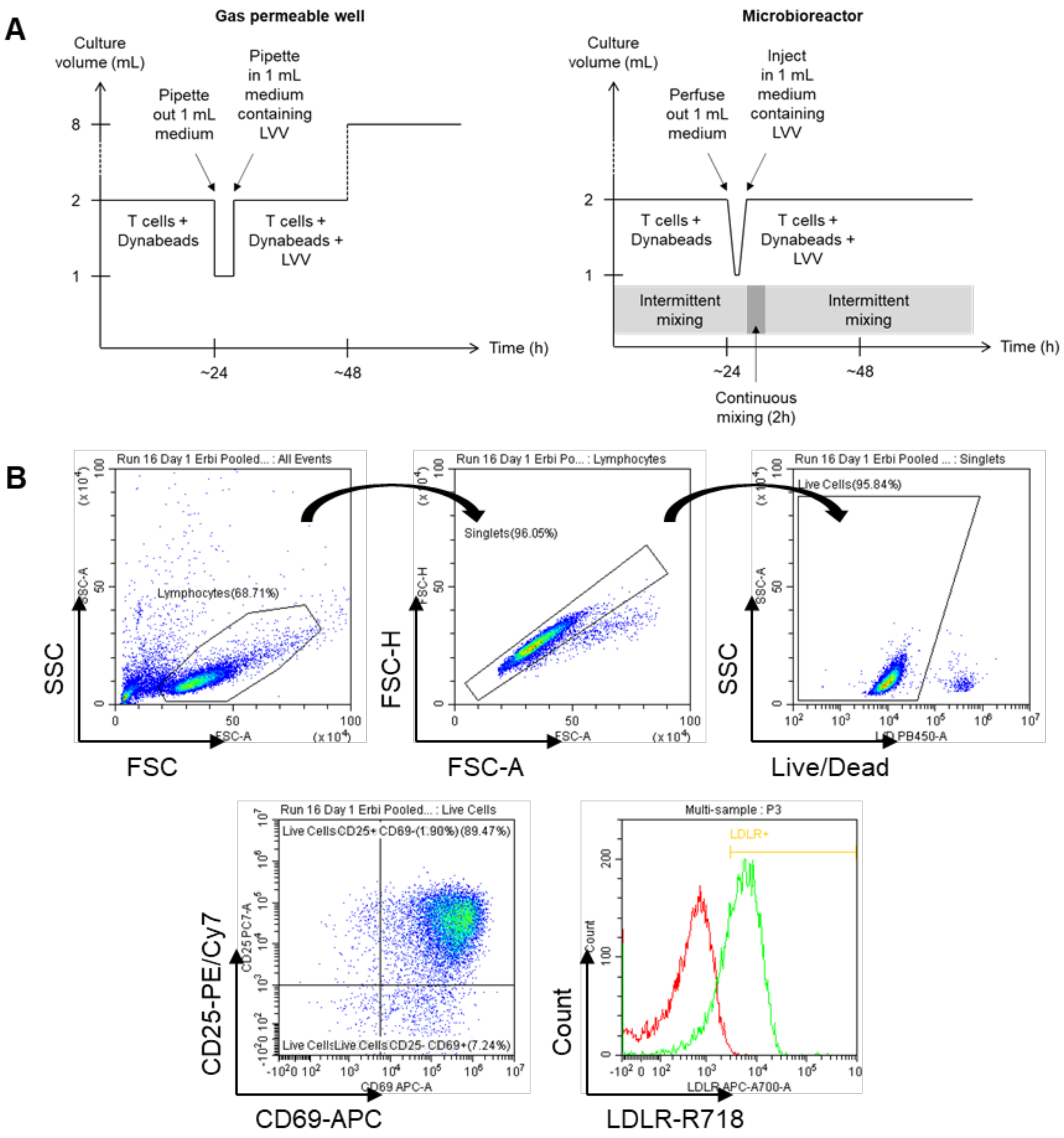


**Supplementary Figure 2: Schematic of activation and transduction process, and gating strategy of T cell activation panel.** (A) Schematic of activation and transduction process. For the gas-permeable wells, 1 mL of cell-free medium was replaced with 1 mL of fresh medium containing diluted LVV, and then left in static culture for 24 h before 6 mL of fresh medium was added to achieve the final culture volume of 8 mL. For the microbioreactor, 1 mL of cell-free medium was removed through the perfusion output, and then 1 mL of fresh medium containing diluted LVV was inoculated through the inoculation port. The growth chamber was continuously mixed for 2 h after the addition of LVV, and then subsequently reverted to intermittent mixing for the remainder of the process. (B) Gating strategy of T cell activation panel. Live, single lymphocytes were gated for CD69 and CD25, as well as LDL-R.

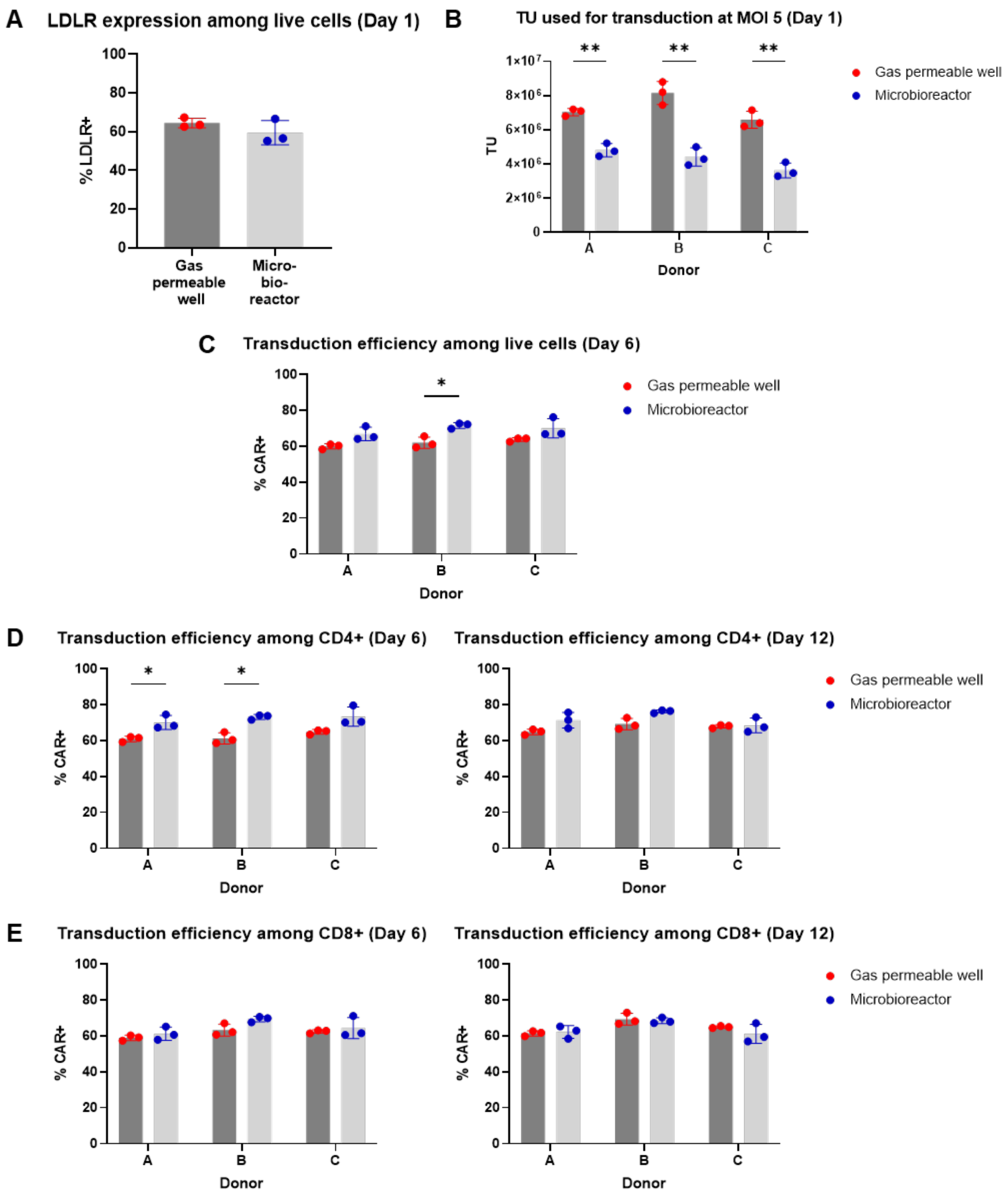


**Supplementary Figure 3: Comparable levels of activation and transduction on the microbioreactor compared with gas-permeable well plate.** (A) Percentage of total live cells expressing LDLR on Day 1. Mean ± SD of n=3 donors. P > 0.05 by unpaired t test. (B) Number of transduction units (TU) of lentiviral vector (LVV) used for transduction on Day 1. Mean ± SD of n=3 wells/cassettes. (C) Percentage of total live cells expressing CAR on Day 6. Mean ± SD of n=3 wells/cassettes. (D) Percentage of CD4+ live cells expressing CAR on Days 6 and 12. Mean ± SD of n=3 wells/cassettes. (E) Percentage of CD8+ live cells expressing CAR on Days 6 and 12. Mean ± SD of n=3 wells/cassettes. (B-E) * $P \leq 0.05$, ** $P \leq 0.01$ by unpaired t test with Holm-Sidak multiple comparisons correction.

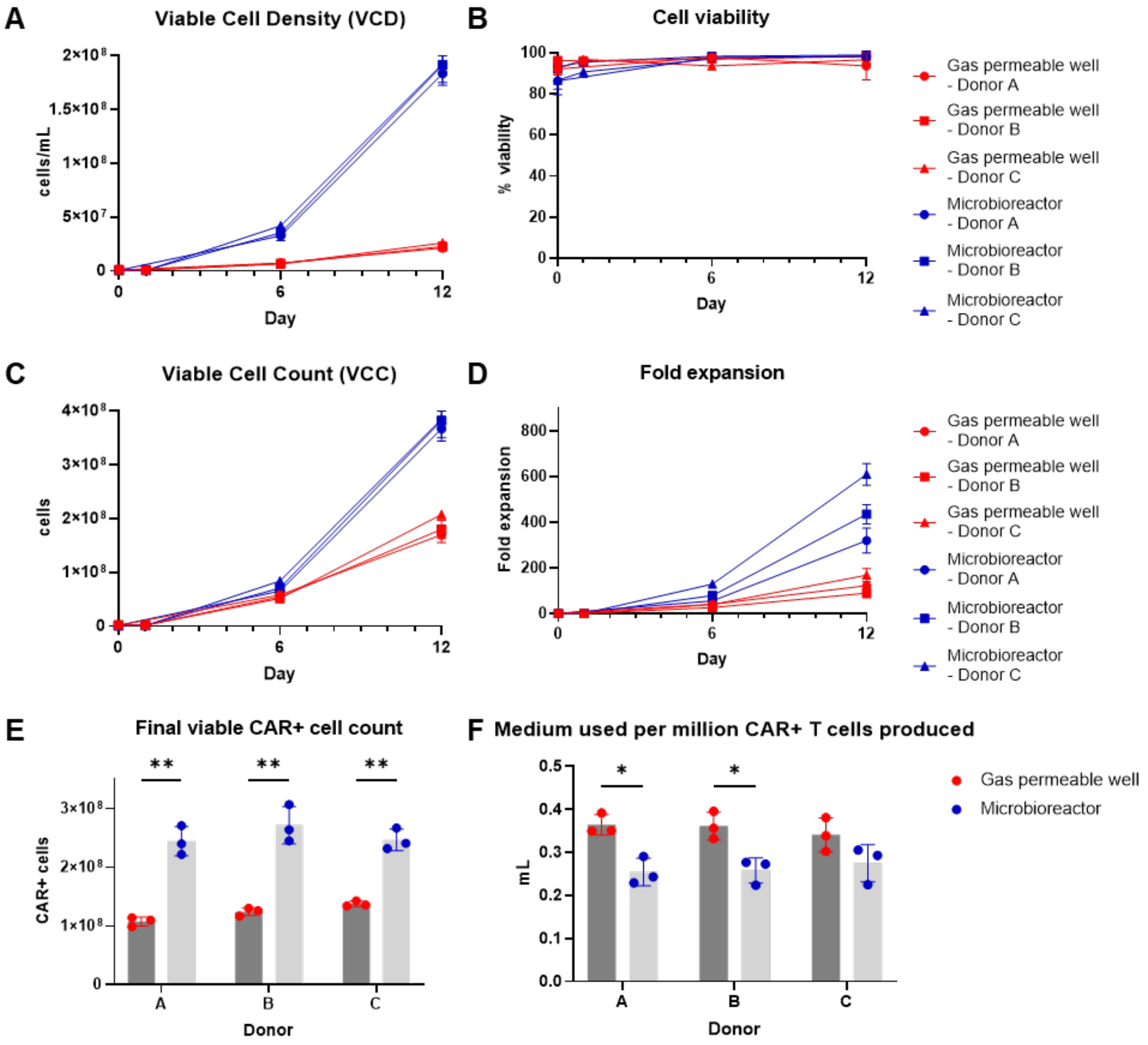


**Supplementary Figure 4: Significantly higher levels of CAR T cell expansion on the microbioreactor compared with gas-permeable well plate.** (A) Viable cell density (VCD) in cells/mL over time, as determined by trypan blue (TB) staining. Mean ± SD of n=3 wells/cassettes. (B) Percentage cell viability as determined by TB staining. Mean ± SD of n=3 wells/cassettes. (C) Viable cell count (VCC) in number of cells over time, as determined by TB staining. Mean ± SD of n=3 wells/cassettes. (D) Fold expansion relative to VCC in cell sample taken after cell inoculation on Day 0, as determined by TB staining. Mean ± SD of n=3 wells/cassettes. (E) Viable CAR+ cell count in number of CAR+ cells on Day 12, as determined by TB staining. Mean ± SD of n=3 wells/cassettes. (F) Volume of medium used per million CAR+ cells produced. Mean ± SD of n=3 wells/cassettes. (E-F) * $P \leq 0.05$, ** $P \leq 0.01$ by unpaired t test with Holm-Sidak multiple comparisons correction.

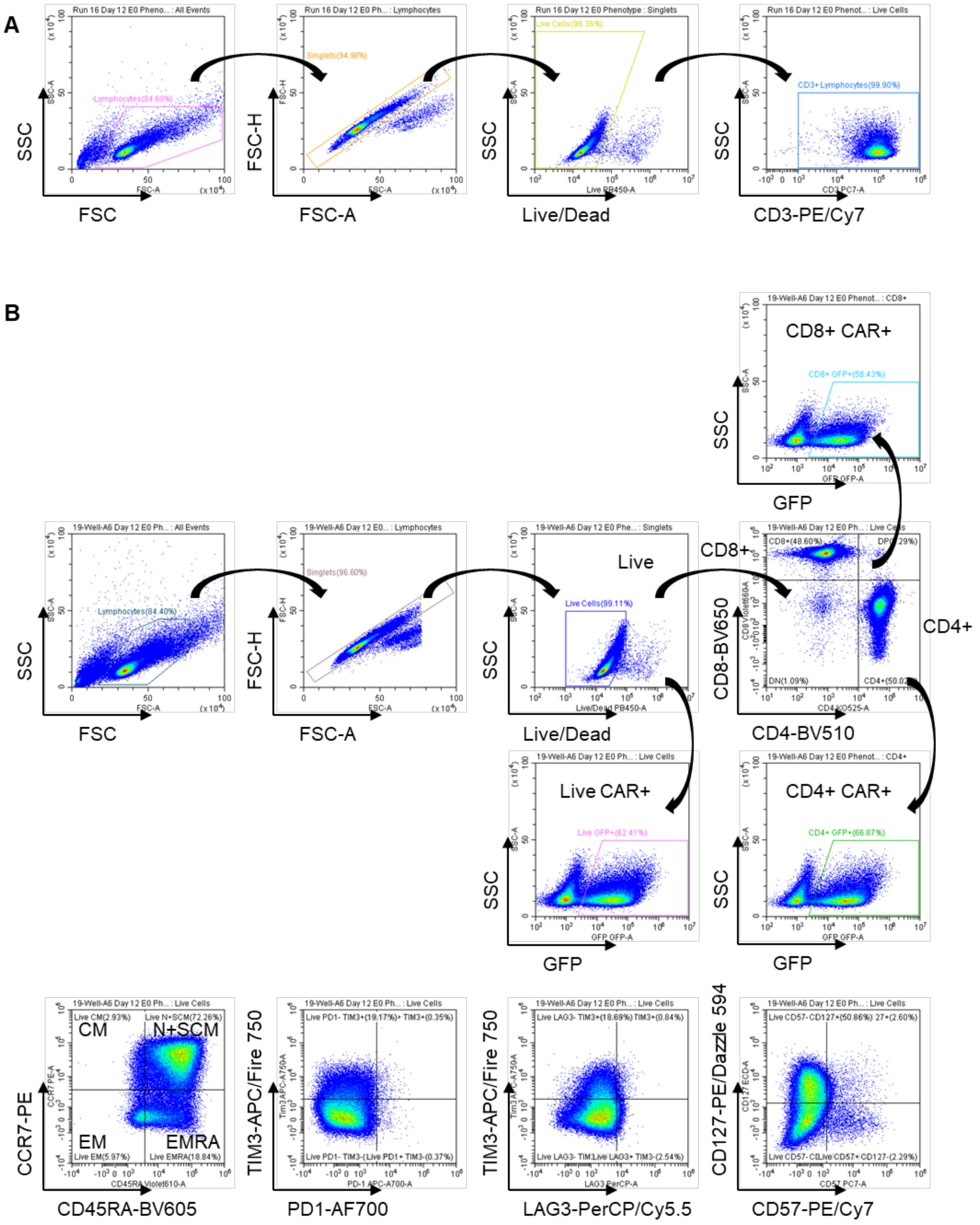


**Supplementary Figure 5: Gating strategy of CAR T cell purity and surface phenotyping panel.** (A) Purity panel: Live, single lymphocytes were gated for CD3. (B) Surface phenotyping panel: Live, single lymphocytes were gated for CD4 and CD8. GFP expression as a proxy of CAR expression was analyzed on Live, CD4+, and CD8+ subsets. Differentiation markers (CCR7 and CD45RA), exhaustion markers (PD1, LAG3, and TIM3), CD127, and CD57 expression were analyzed on Live, Live CAR+, CD4+, CD4 CAR+, CD8+, and CD8+ CAR+ subsets.

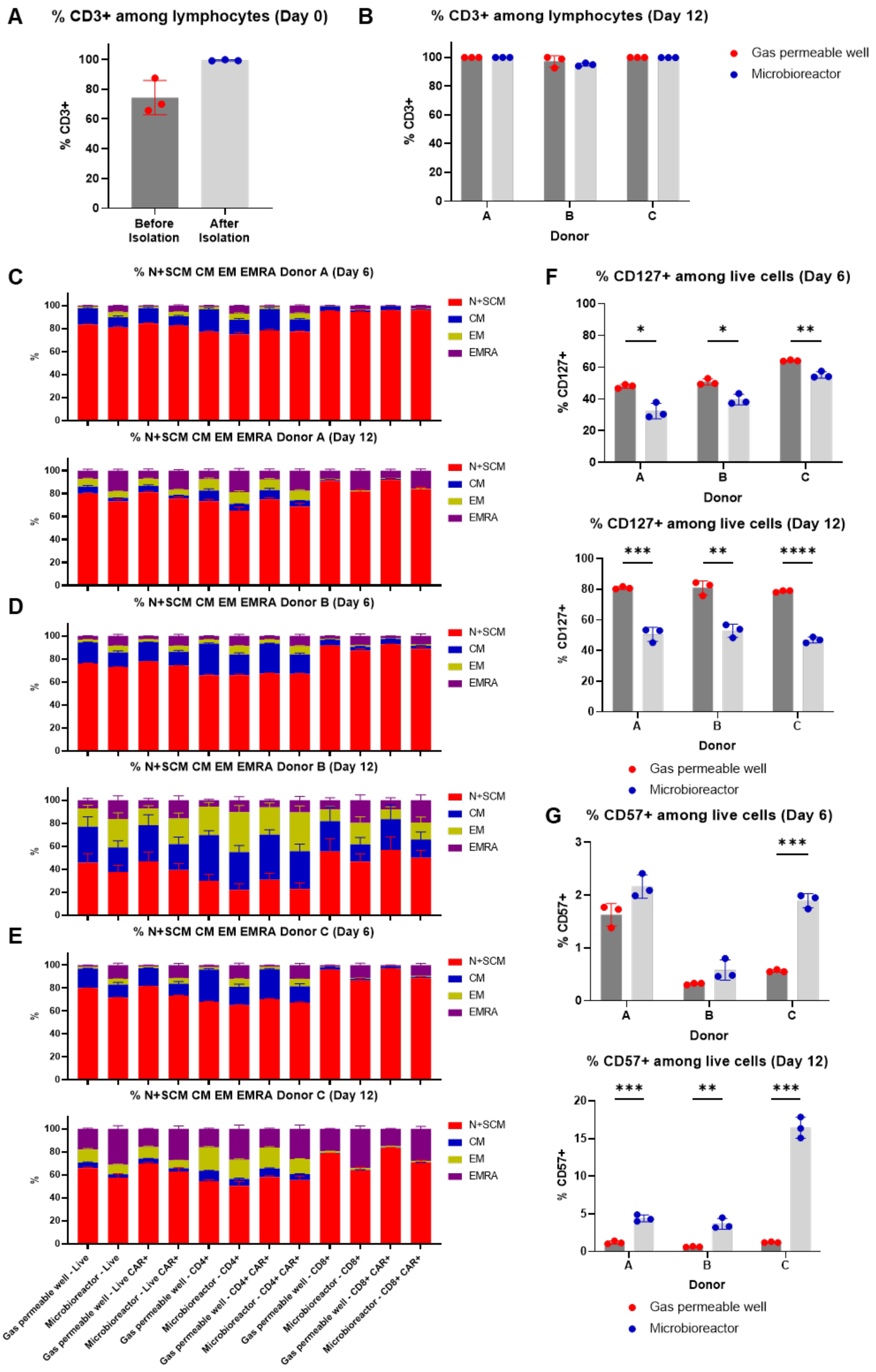
A
% CD3+ among lymphocytes (Day 0)
% CD3+
Before Isolation
After Isolation
B
% CD3+ among lymphocytes (Day 12)
Gas permeable well
Microbioreactor
Donor
C
% N+SCM CM EM EMRA Donor A (Day 6)
% N+SCM CM EM EMRA Donor A (Day 12)
D
% N+SCM CM EM EMRA Donor B (Day 6)
% N+SCM CM EM EMRA Donor B (Day 12)
E
% N+SCM CM EM EMRA Donor C (Day 6)
% N+SCM CM EM EMRA Donor C (Day 12)
N+SCM
CM
EM
EMRA
Gas permeable well - Live
Microbioreactor - Live
Gas permeable well - Live CAR+
Microbioreactor - Live CAR+
Gas permeable well - CD4+
Microbioreactor - CD4+
Gas permeable well - CD4+ CAR+
Microbioreactor - CD4+ CAR+
Gas permeable well - CD8+
Microbioreactor - CD8+
Gas permeable well - CD8+ CAR+
Microbioreactor - CD8+ CAR+
F
% CD127+ among live cells (Day 6)
% CD127+ among live cells (Day 12)
% CD127+
G
% CD57+ among live cells (Day 6)
% CD57+ among live cells (Day 12)
% CD57+
Donor
Gas permeable well
Microbioreactor

**Supplementary Figure 6: T cell differentiation and exhaustion phenotypes of CAR T cells from the microbioreactor and the gas-permeable well plate.** (A) Percentage of CD3+ T cells (gated on lymphocytes) before and after T cell isolation on Day 0. Mean ± SD of n=3 donors. (B) Percentage of CD3+ T cells (gated on lymphocytes) on Day 12. Mean ± SD of n=3 wells/cassettes. (C-E) Percentage of CCR7+ CD45RA+ (naïve and stem cell memory, Tn/Tscm), CCR7+ CD45RA– (central memory, Tcm), CCR7– CD45RA– (effector memory, Tem), and CCR7– CD45RA+ (terminally differentiated effector, Temra) cells gated on total live cells ("Live"), on CAR+ live cells ("Live CAR+"), on CD4+ cells ("CD4+"), on CAR+ CD4+ cells ("CD4+ CAR+"), on CD8+ cells ("CD8+"), and on CAR+ CD8+ cells ("CD8+ CAR+") on Days 6 and 12 for Donor A, B, and C, respectively. Mean ± SD of n=3 wells/cassettes. (F-G) Percentage of total live cells expressing CD127 (F) and CD57 (G) on Days 6 and 12. Mean ± SD of n=3 wells/cassettes. * $P \leq 0.05$, ** $P \leq 0.01$, *** $P \leq 0.001$, **** $P \leq 0.0001$ by unpaired t test with Holm-Sidak's multiple comparisons correction.

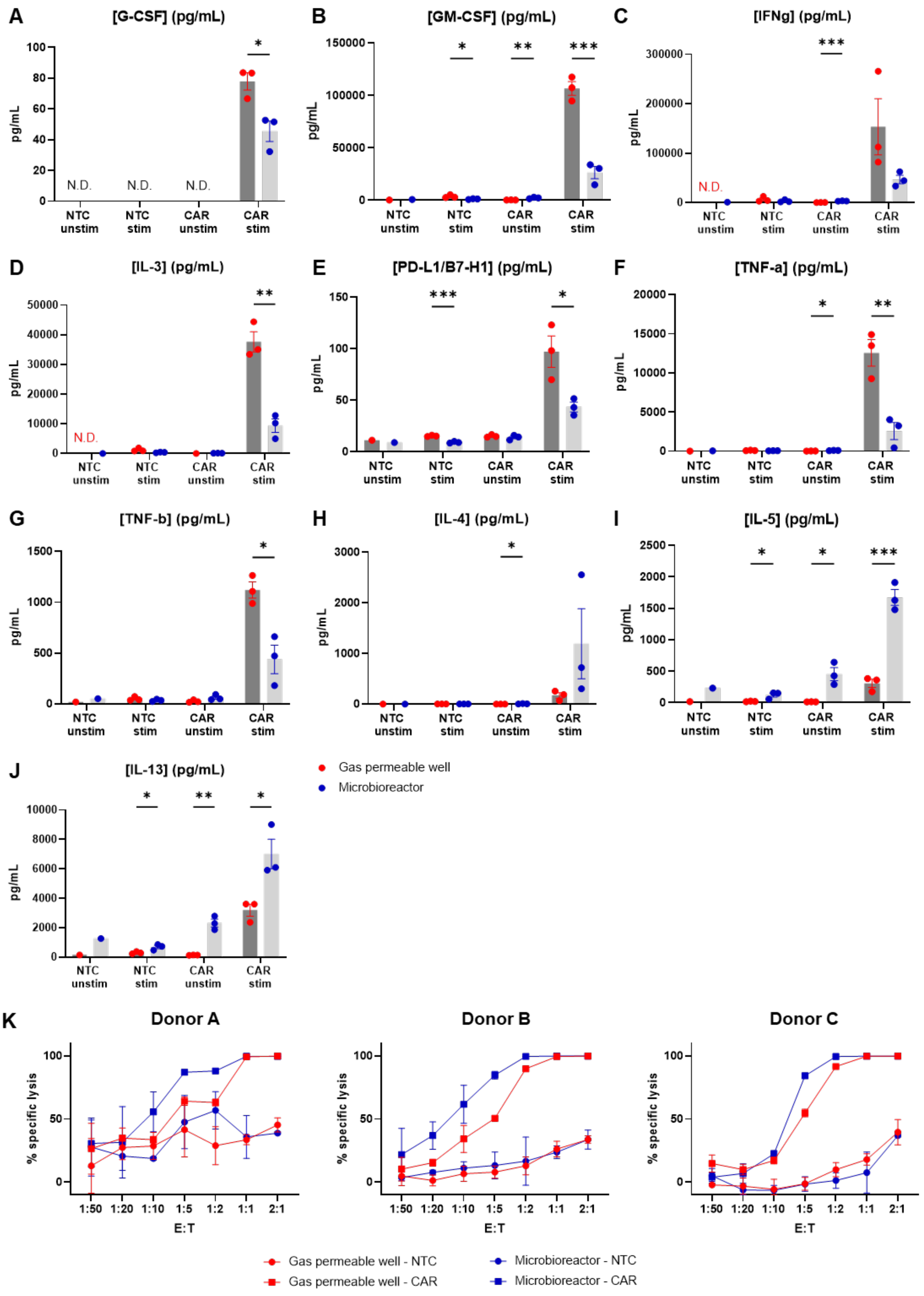
A
[G-CSF] (pg/mL)
B
[GM-CSF] (pg/mL)
C
[IFNg] (pg/mL)
D
[IL-3] (pg/mL)
E
[PD-L1/B7-H1] (pg/mL)
F
[TNF-a] (pg/mL)
G
[TNF-b] (pg/mL)
H
[IL-4] (pg/mL)
I
[IL-5] (pg/mL)
J
[IL-13] (pg/mL)
pg/mL
N.D.
NTC unstim
NTC stim
CAR unstim
CAR stim
Gas permeable well
Microbioreactor
K
Donor A
Donor B
Donor C
% specific lysis
E:T
1:50
1:20
1:10
1:5
1:2
1:1
2:1
Gas permeable well - NTC
Gas permeable well - CAR
Microbioreactor - NTC
Microbioreactor - CAR

**Supplementary Figure 7: CAR T cells produced on the microbioreactor were highly functional in terms of cytokine release and cytolytic activity against tumor cells.** (A-J) Cytokine secretion following overnight co-culture of T cells and NALM6 cells at a 1:1 CAR+ T cell:NALM6 cell ratio. Cell culture supernatant from the co-culture of cells from one technical replicate (one well/cassette) of each donor were used for this assay. Mean ± SEM of n=3 donors (except for NTC unstim, for which only Donor A was measured), each assayed in technical triplicates. NTC: non-transduced control. CAR: CAR T cells (unsorted). Unstim: T cells alone. Stim: Co-cultured with NALM6 cells. * $P \leq 0.05$, ** $P \leq 0.01$, *** $P \leq 0.001$ by unpaired t test with Holm-Sidak's multiple comparisons correction. (K) Percentage specific lysis of NALM6 cells following overnight co-culture of T cells and NALM6 cells at various CAR+ T cell:NALM6 cell ratios. Cryopreserved cells from one technical replicate (one well/cassette) of each donor were thawed for this assay. Percentage specific lysis was determined by flow cytometry and calculated relative to wells with NALM6 alone (0% lysis). Mean ± SD of n=3 technical replicates.

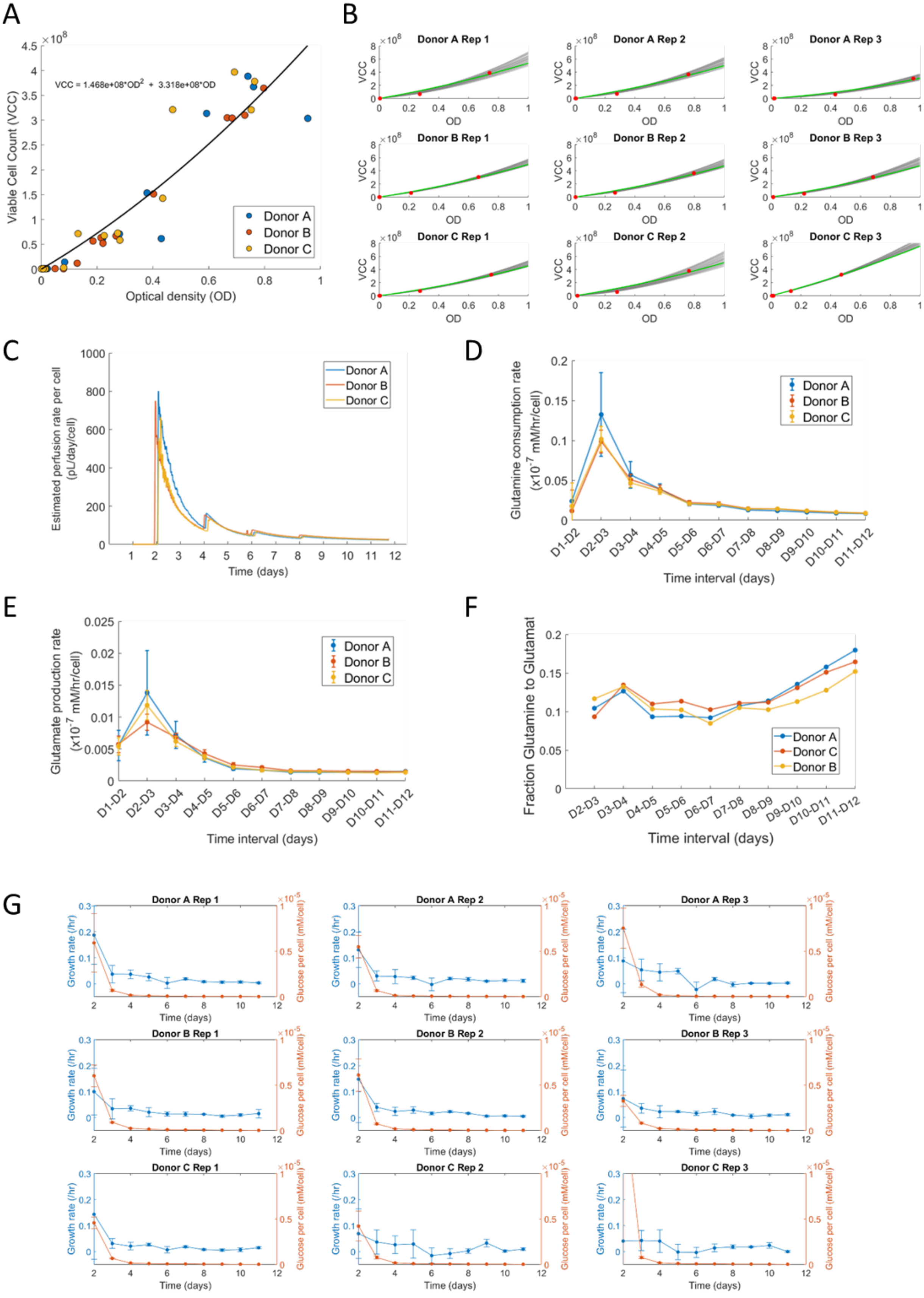
A
Viable Cell Count (VCC)
Optical density (OD)
VCC = 1.468e+08*OD^2 + 3.318e+08*OD
Donor A
Donor B
Donor C
B
Donor A Rep 1
Donor A Rep 2
Donor A Rep 3
Donor B Rep 1
Donor B Rep 2
Donor B Rep 3
Donor C Rep 1
Donor C Rep 2
Donor C Rep 3
VCC
OD
C
Estimated perfusion rate per cell (pL/day/cell)
Time (days)
D
Glutamine consumption rate (x10^-7 mM/hr/cell)
Time interval (days)
E
Glutamate production rate (x10^-7 mM/hr/cell)
Time interval (days)
F
Fraction Glutamine to Glutamat
Time interval (days)
Donor A
Donor C
Donor B
G
Growth rate (/hr)
Glucose per cell (mM/cell)
Time (days)

**Supplementary Figure 8: Estimating growth rates and metabolic rates using computational modeling.** (A) The generalized OD-VCC fit equation. Scatter plots show the measured cell count values and the corresponding OD values obtained from the bioreactor (color coded by donor). The fitted curve is in black and the fit equation is displayed within the plot. (B) Family-of-fit curves (set of customized OD-VCC curves) for each donor and replicate. Red points indicate the measured cell counts and corresponding OD values. Green curve is the linear-customized fit (See Methods), and the grey curves are the individual curves that make up the family-of-fits for the corresponding donor/replicate. (C) Estimated perfusion rate per cell for the three donors. Cell numbers were estimated from OD values, using 50 randomly chosen fit equations from the family-of-fits we originally established for each donor and replicate. Plotted are the mean values of perfusion rate/cell obtained from 150 values (3 replicates*50 equations for each replicate). (D) Plots showing the estimated 24-hour average glutamine consumption rates ($x10^{-7}$ mM glutamine/cell/hr) for the three donors from Days 1-12. Model-based estimation used the workflow in **Figure 6A** and 50 different OD-VCC fit equations per replicate (chosen at random from the family-of-fits for that donor/replicate). The x-axis shows 24-hour time intervals (Day1-Day2, Day2-Day3, etc). (E) Same as (D) but for glutamate production rate ($x10^{-7}$ mM glutamate/cell/hr). (F) Plot showing the apparent fraction of glutamine funneled towards glutamate for the three donors from Days 1-12. This is computed as the model-estimated mean glutamate production rate divided by the model-estimated mean glutamine consumption rate. (G) Plots showing the measured glucose concentration per cell in the bioreactor (orange) and the average estimated growth rates at the same timepoints as glucose measurements (blue) for each donor/replicate. The y-axis for the glucose concentration per cell (orange) for all panels has been truncated at an upper bound of $1x10^{-5}$ mM/cell for visual clarity. This omits the mean glucose concentration per cell for Donor C Replicate 3 at Day 2, which was $1.849x10^{-5}$ mM/cell.

A

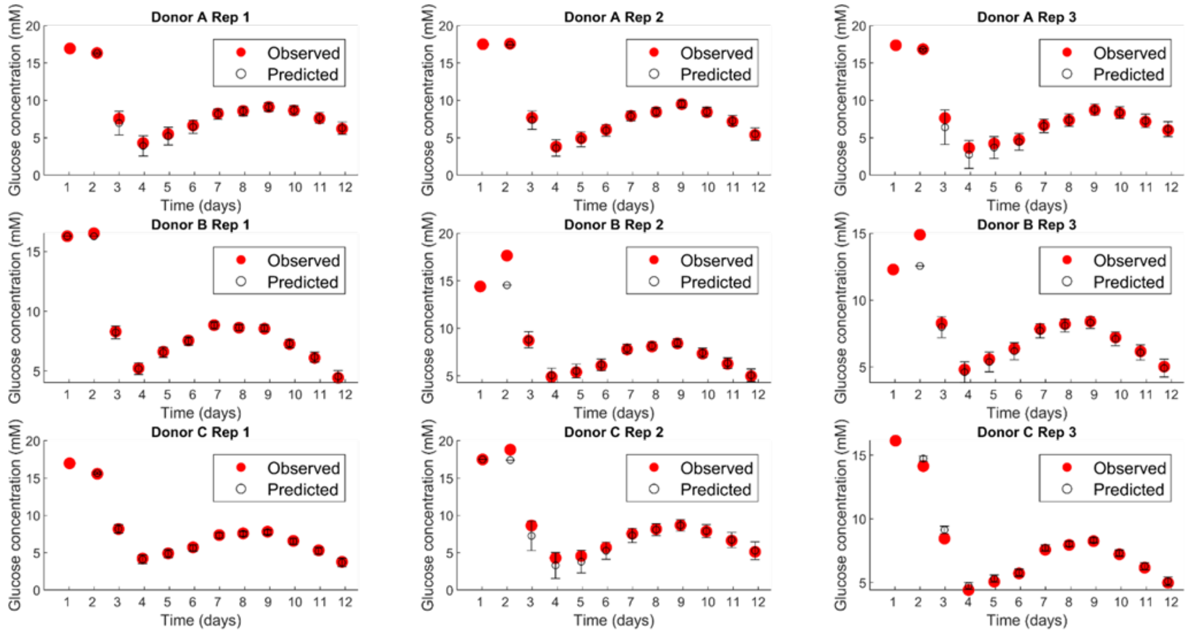


B

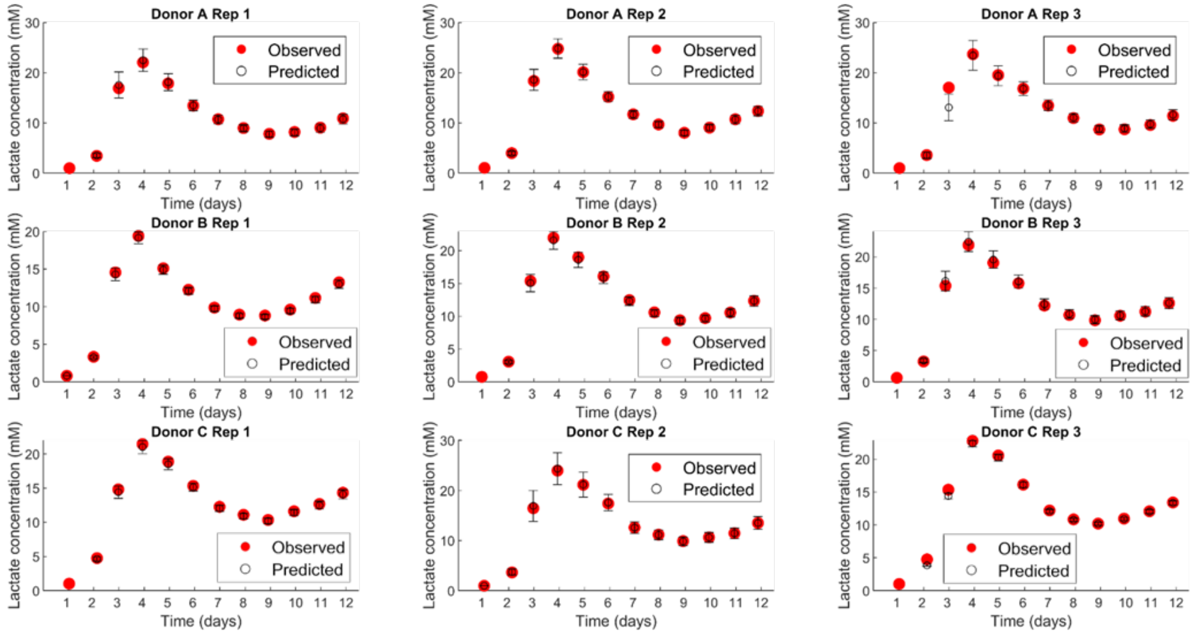


**Supplementary Figure 9: Quality of fit between model-estimated metabolite concentrations and measured metabolite concentrations for Glucose and Lactate.** (A) Experimentally measured (red) vs model-estimated average glucose concentration (black). Estimated concentration is computed using ordinary differential equations, into which we plugged in mean glucose consumption rates for each day and donor (from **Figure 6D**), and cell numbers estimated from OD (using 50 randomly chosen fit equations from the family-of-fits we originally established for each donor and replicate). (B) Same as (A) but for concentrations of lactate.

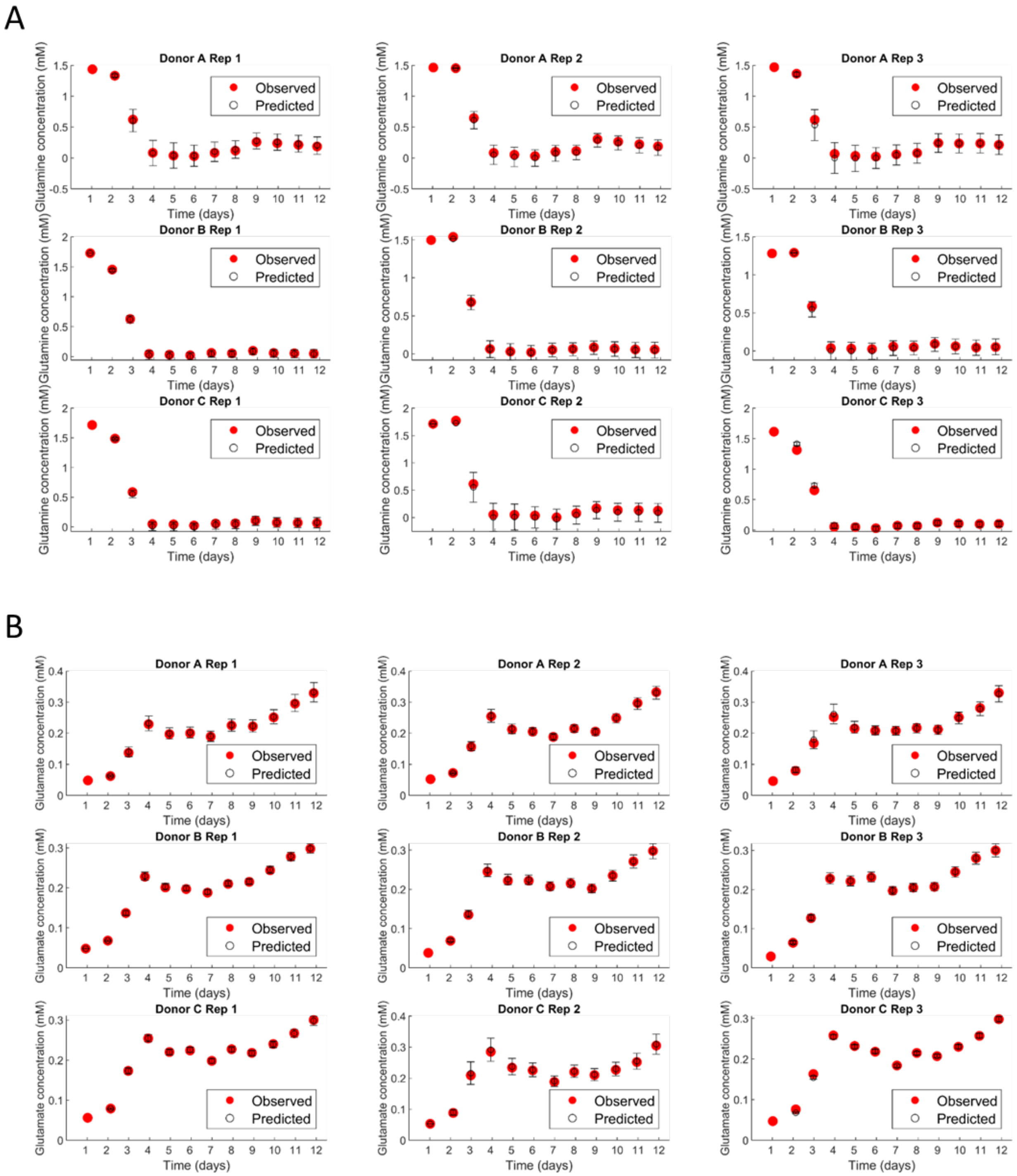


**Supplementary Figure 10: Quality of fit between model-estimated metabolite concentrations and measured metabolite concentrations for Glutamine and Glutamate.** (A) Experimentally measured (red) vs model-estimated average glutamine concentration (black). Estimated concentration is computed using ordinary differential equations, into which we plugged in mean glutamine consumption rates for each day and donor (from **Supplementary Figure 8D**), and cell numbers estimated from OD (using 50 randomly chosen fit equations from the family-of-fits we originally established for each donor and replicate). (B) Same as (A) but for concentrations of glutamate.

**Supplementary Table 1: Exact timepoints of perfusion rate increases for each donor and replicate.**

| Donor | Perfusion rate increased to (vvd) | Replicate 1 (hr) | Replicate 2 (hr) | Replicate 3 (hr) | Row mean (hr) |
|---|---|---|---|---|---|
| A | 1 | 49.55 | 49.56 | 49.49 | 49.54 |
| | 2 | 97.74 | 97.73 | 97.66 | 97.71 |
| | 3 | 145.83 | 145.85 | 145.75 | 145.81 |
| | 4 | 193.83 | 193.85 | 193.75 | 193.81 |
| B | 1 | 46.35 | 46.31 | 46.28 | 46.31 |
| | 2 | 95.95 | 95.91 | 95.88 | 95.91 |
| | 3 | 143.82 | 143.81 | 143.78 | 143.80 |
| | 4 | 191.84 | 191.82 | 191.81 | 191.82 |
| C | 1 | 49.64 | 49.60 | 49.54 | 49.59 |
| | 2 | 99.74 | 99.70 | 99.64 | 99.69 |
| | 3 | 147.64 | 147.60 | 147.54 | 147.59 |
| | 4 | 195.59 | 195.55 | 195.49 | 195.54 |

**Supplementary Table 2: Growth rates, glucose and glutamine consumption rates, lactate and glutamate production rates, and oxygen consumption rates of CAR T cells.** Growth rates and oxygen consumption rates are continuous (instantaneous) estimates, but for robustness, we computed peak and final values by averaging over 3-hr windows. Specifically, peak values are computed from a 3-hr window around the highest mean rate for each donor, and the final rate is computed from a 3-hr window around the timepoint t = 276hr (= 11.5 days). Metabolite consumption/production rates are available for each 24-hr interval, so we took peak values from the highest estimated 24-hr average rate and final rates from the time interval Day11-Day12.

| | **Donor A (mean ± S.D)** | **Donor B (mean ± S.D)** | **Donor C (mean ± S.D)** |
|---|---|---|---|
| Peak Growth rate ($hr^{-1}$) | 0.1012 ± 0.0444 | 0.0837 ± 0.0376 | 0.1186 ± 0.1025 |
| Final Growth rate ($hr^{-1}$) | 0.0041 ± 0.0014 | 0.0052 ± 0.0017 | 0.0038 ± 0.0036 |
| Peak oxygen consumption rate (mM/hr/cell) | 0.148e-4 ± 0.052e-4 | 0.219e-4 ± 0.108e-4 | 0.89e-4 ± 1.23e-4 |
| Final Oxygen consumption rate (mM/hr/cell) | 0.438e-6 ± 0.123e-6 | 0.64e-6 ± 0.099e-6 | 0.689e-6 ± 0.147e-6 |
| Peak Glucose consumption rate ($\times 10^{-7}$ mM/hr/cell) | 1.2626 ± 0.475 | 0.8201 ± 0.1195 | 0.7597 ± 0.1311 |
| Final Glucose consumption rate ($\times 10^{-7}$ mM/hr/cell) | 0.0533 ± 0.0074 | 0.0610 ± 0.0072 | 0.0612 ± 0.0069 |
| Peak Glutamine consumption rate ($\times 10^{-7}$ mM/hr/cell) | 0.1324 ± 0.0525 | 0.0985 ± 0.0143 | 0.1016 ± 0.0158 |
| Final Glutamine consumption rate ($\times 10^{-7}$ mM/hr/cell) | 0.0083 ± 0.0011 | 0.0087 ± 9.18e-4 | 0.0089 ± 7.81e-4 |
| Peak Lactate production rate ($\times 10^{-7}$ mM/hr/cell) | 1.7946 ± 0.26 | 1.4123 ± 0.1621 | 1.2554 ± 0.2149 |
| Final Lactate production rate ($\times 10^{-7}$ mM/hr/cell) | 0.0572 ± 0.0084 | 0.0678 ± 0.0083 | 0.0695 ± 0.0069 |
| Peak Glutamate production rate ($\times 10^{-7}$ mM/hr/cell) | 0.0138 ± 0.0066 | 0.0092 ± 0.0013 | 0.0118 ± 0.0023 |
| Final Glutamate production rate ($\times 10^{-7}$ mM/hr/cell) | 0.0015 ± 1.99e-4 | 0.0014 ± 1.56e-4 | 0.0014 ± 1.11e-4 |
